# A facile method of mapping HIV-1 neutralizing epitopes using chemically masked cysteines and deep sequencing

**DOI:** 10.1101/2020.04.15.041996

**Authors:** Rohini Datta, Rohan Roy Chowdhury, Kavyashree Manjunath, Luke Elizabeth Hanna, Raghavan Varadarajan

## Abstract

Identification of specific epitopes targeted by neutralizing antibodies is essential to advance epitope-based vaccine design strategies. We report a facile methodology for rapid epitope mapping of neutralizing antibodies (NAbs) against HIV-1 Envelope (Env) at single residue resolution, using Cys labeling, viral neutralization assays and deep sequencing. This was achieved by the generation of a library of Cys mutations in Env glycoprotein on the viral surface, covalent labeling of the Cys residues using a Cys reactive label that masks epitope residues, followed by infection of the labeled mutant virions in mammalian cells in the presence of NAbs. Env gene sequencing from NAb-resistant viruses was used to accurately delineate epitopes for the NAbs VRC01, PGT128 and PGT151. These agreed well with corresponding experimentally determined structural epitopes previously inferred from NAb:Env structures. HIV-1 infection is associated with complex and polyclonal antibody responses, typically composed of multiple antibody specificities. Deconvoluting the epitope specificities in a polyclonal response is a challenging task. We therefore extended our methodology to map multiple specificities of epitopes targeted in polyclonal sera, elicited in immunized animals as well as in an HIV-1 infected elite neutralizer capable of neutralizing Tier-3 pseudoviruses with high titers. The method can be readily extended to other viruses for which convenient reverse genetics or lentiviral surface display systems are available.

## Introduction

The primary objective of a vaccine against a viral infection is to elicit a long-lasting protective neutralizing antibody response. The design of an effective vaccine candidate necessitates a high-resolution, residue-level, map of the epitopes on the viral surface that are targeted by neutralizing antibodies (NAbs). This information is usually obtained from X-ray crystallography or cryo-EM based structural studies of purified complexes of a monoclonal antibody with its target viral protein. These methods provide a detailed high-resolution map of the interaction between the antibody and its target surface and assist in vaccine design. However, apart from being highly labor-intensive and time-consuming, these methods are not suitable for mapping polyclonal antibody responses during a viral infection. Therefore, there is an unmet requirement for a rapid, parallelizable, complementary method to accurately map polyclonal neutralizing antibody epitopes. We recently described a methodology to decipher epitopes of monoclonal antibody panels using yeast surface display, Cys labeling and deep sequencing (1, 2). Here we have combined Cys labeling of viral surface glycoprotein and deep sequencing, with virus neutralization assays to map epitopes of neutralizing monoclonal antibodies and polyclonal sera against the HIV-1 virus. The design of an effective vaccine against HIV-1 has met with limited success so far, because of the inability of immunogens to elicit broadly neutralizing antibodies (bNAbs) targeted to the trimeric envelope (Env) glycoprotein, the sole viral antigen on the surface of the virus (3–5). Most bNAbs isolated from infected individuals are directed to one of the major sites of vulnerability on Env: V1/V2 loop apex, V3 loop, CD4-binding site, center of the gp120 silent face, gp120-gp41 subunit interface, gp41 fusion peptide and membrane proximal external region (MPER) of gp41 (3, 6, 7).

We developed a high-throughput methodology for mapping neutralizing HIV-1 epitopes at single residue resolution directly on the native viral Env, obviating the need to purify Env glycoprotein. To this end, a library of single-site Cys mutations were engineered at selected solvent-exposed sites in the Env glycoprotein on the surface of the virus. Cys mutants were covalently labeled using a Cys-reactive bulky maleimide label, that masks epitope residues, followed by infection of the labeled mutant viruses in mammalian cells in the presence of NAbs. Subsequently, deep sequencing of the env gene from NAb-resistant viruses was used to delineate epitopes of NAbs (Figure 1). We focused our epitope mapping efforts to the Env ectodomain of the clade B, CCR5-tropic primary HIV-1 isolate JRFL, which has been extensively characterized both biochemically and structurally, and is used as a model primary virus.

**Figure 1.**
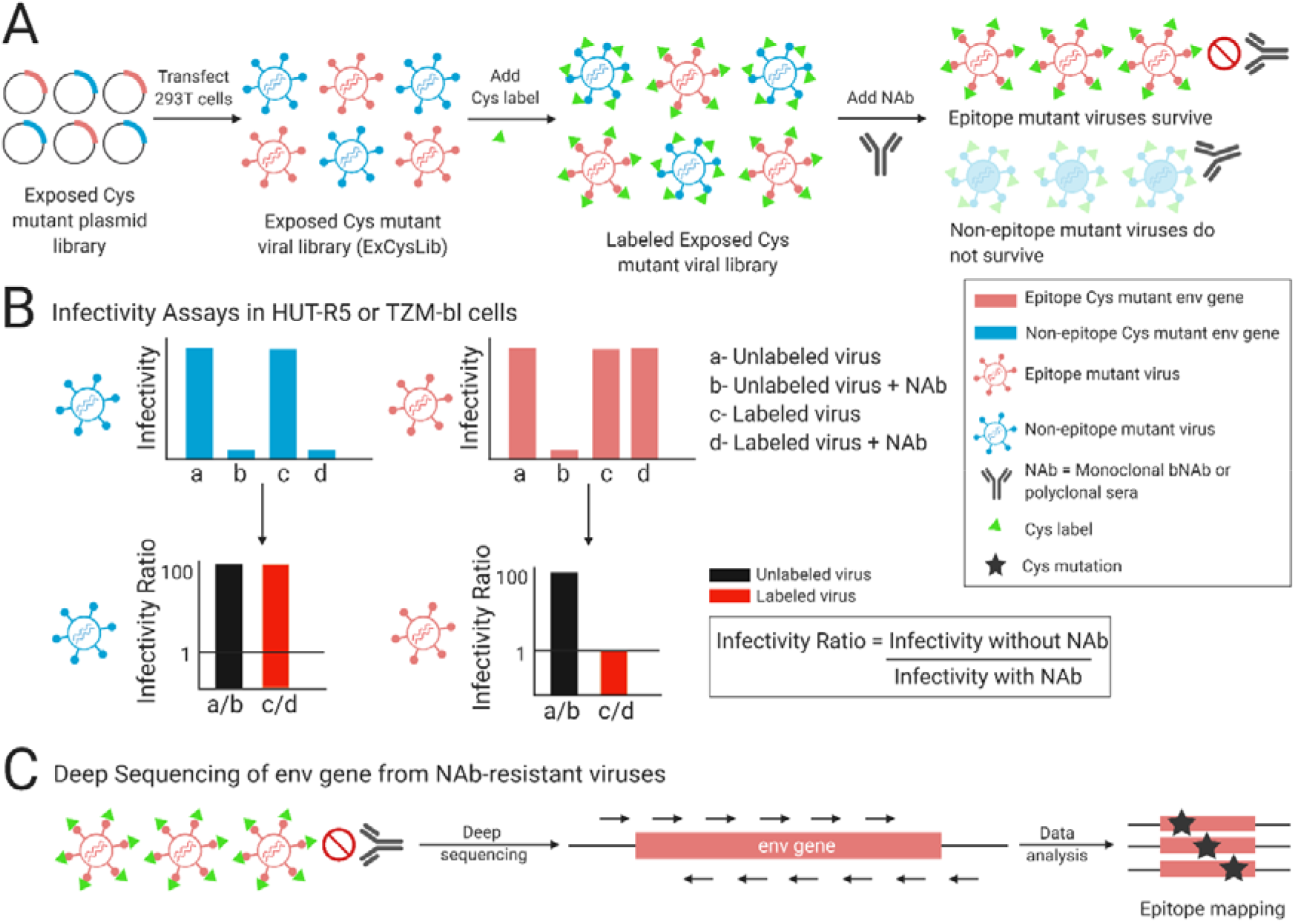
Approach to rapidly map neutralizing epitopes of HIV-1. (A) A library of single Cys mutants was generated at 81 solvent exposed sites in the env gene of the HIV-1 molecular clone, pLAI-JRFL. Approximately half of these sites are part of known bNAb epitopes, whereas the rest are non-epitopic. The exposed Cys mutant plasmid library, an equimolar pool of these single Cys env mutants in pLAI-JRFL, was used to transfect HEK293T cells. This led to the production of a library of mutant viruses, having Cys mutations at the selected solvent exposed residues in Env; henceforth referred to as the ExCysLib. The ExCysLib was covalently labeled with a Cys label, EZ-Link™ Maleimide-PEG2-Biotin, and then incubated with NAbs (either monoclonal bNAbs or polyclonal sera). At epitope residues, masking by the bulky Cys label prevents bNAbs from binding and neutralizing the epitope mutant viruses. (B) Quantification using infectivity assays of the ExCysLib before and after labeling, in the absence and presence of NAbs. Unlabeled epitope and non-epitope Cys mutant viruses are similarly sensitive to neutralization. However, upon labeling, epitope mutant viruses are resistant to neutralization, unlike the non-epitope counterparts. Infectivity ratio (= infectivity without NAb/infectivity with NAb) was used as a metric to discriminate between epitope versus non-epitope mutant viruses. Upon labeling, infectivity ratio of non-epitope mutant viruses >> 1, however for epitope mutant viruses, the ratio is ~1. (C) Deep sequencing of the env gene from antibody/sera-resistant virus pools was used to identify epitope residues at single residue resolution. The figure was created using Bio-Render.com

## Results

### Residue selection and generation of ExCysLib

A total of 81 solvent-exposed residues (Table S1) were selected from the X-ray crystal structure of the pre-fusion HIV-1 Env ectodomain using a combined criterion of >30% solvent accessibility and centroid distance between residue side-chains >8 Å (Figure S1). The side-chain side-chain centroid distance cut-off was chosen to discard residues in close proximity (8), which are likely to be part of the same Ab epitope (9). This led to the identification of a set of solvent-exposed residues which collectively exhibit uniform surface coverage of the Env trimer (Figure S2). 47 out of the 81 selected residues are epitopes of known bNAbs or contact sites for CD4, the primary cellular receptor of HIV. Cys mutations were individually engineered at the selected sites in the env gene in the pLAI-JRFL molecular clone. This molecular clone encodes the full-length infectious genome of the LAI-JRFL virus. The Env ectodomain in this molecular clone is from the JRFL strain (residues 25 – 690), whereas the rest of the sequence is derived from the LAI strain of HIV-1 (10). Cys mutants in pLAI-JRFL were pooled in equimolar quantities to create a mutant plasmid DNA library, which was transfected in HEK293T cells to produce the Exposed Cys mutant viral Library (ExCysLib) (Figure 1A), a library of single-site Cys mutant viruses. In single-cycle and multi-cycle infectivity assays, the ExCysLib retained significant viral infectivity (Figure S3) and was therefore used for epitope mapping studies.

### Significant viral infectivity is retained after Cys labeling of Env

Viral supernatants from HEK293T cells contain 20% Fetal Bovine Serum (FBS) which has a high protein content (~30-40 mg/ml) and includes a large number of sulfhydryl groups that can be labeled. This would necessitate the use of high concentrations of the Cys label for our experiments. In order to monitor the labeling kinetics of Cys residues under these conditions, we exploited the strong interaction between two model proteins, CcdB and DNA Gyrase (8) to quantify the extent of Cys labeling in the presence of 20% FBS, as a proxy for viral labeling (Figure S4A). CcdB is a bacterial toxin which binds with high affinity to the GyrA subunit of its cellular target DNA Gyrase, as well as to the GyrA14 fragment that consists of residues 363-494 of GyrA (11). In the present work we employed a GyrA14 mutant with a Cys residue engineered at a solvent exposed site (I491C) that lies outside the CcdB binding site. GyrA14-I491C was labeled using different concentrations of the Cys labeling reagent, EZ-Link™ Maleimide-PEG2-Biotin, for different time points of incubation, in the presence of 20% FBS to mimic viral labeling. The labeled GyrA14-I491C was then purified using a CcdB affinity column, and mass of eluted protein was analyzed using MALDI-TOF mass spectrometry (Figure S4A) to determine labeling kinetics. Labeling was maximal at a concentration of 10 mM labeling reagent and largely complete after two hours of incubation (Figure S4B).

Next, we incubated the WT virus with 1 and 10 mM of the Cys label for two hours and measured single-cycle viral infectivity in TZM-bl cells. Since the use of 10 mM label likely led to a high extent of label incorporation without significantly affecting viral infectivity (Figure S5), we used this concentration in all subsequent experiments.

### ExCysLib can be used to map distinct bNAb epitopes

To validate our methodology, proof-of-concept studies were carried out with the ExCysLib to map epitopes of bNAbs directed to distinct sites on Env. To decide the amount of antibody to use, we performed neutralization assays of WT LAI-JRFL virus in TZM-bl cells, in the presence of the following bNAbs: CD4-binding site directed VRC01, V3-glycan targeting PGT128 and gp120-gp41 interface-specific PGT151. For epitope mapping studies, we used bNAb concentrations 100 times in excess of the measured IC_50_ with LAI-JRFL (Table S2) (12, 13).

Approximately half of the residues selected for the ExCysLib are part of known bNAb epitopes. The ExCysLib, a mixed pool of Cys mutants within and outside the epitope for a given bNAb, was covalently labeled with a Cys label, and then incubated with NAbs (monoclonal bNAbs or polyclonal sera) (Figure 1A). Upon labeling, masking by the bulky Cys label should prevent NAbs from neutralizing those mutant viruses that contain the Cys residue within the epitope. However, mutant viruses that contain the Cys residue outside the epitope will continue to be sensitive to neutralization, even after labeling (Figure 1B). Thus, labeled mutant viruses that contain the introduced Cys within or outside the epitope can be distinguished using the infectivity ratio (=infectivity without NAb/infectivity with NAb). These will have infectivity ratios ~1 or >>1 respectively (Figure 1B).

Mapping NAb epitopes using the entire ExCysLib necessitates deep sequencing of env gene from NAb-resistant virion pools (Figure 1C). This was accomplished using HUT-R5, a T-cell line permissive to multi-cycle infection (14). To validate our methodology, we first mapped known bNAb epitopes. For this, we carried out multicycle infectivity assays of the labeled ExCysLib in the presence of 100 IC_50_ equivalents of bNAbs VRC01, PGT128 or PGT151. In multi-cycle infectivity assays, the cells are split every alternate day post-infection. Cys labeling is carried out at each time point after splitting, by adding labeling reagent to maintain a total concentration of 10 mM Maleimide-PEG2-Biotin. The label is added two hours before adding additional neutralizing antibody to maintain a total antibody concentration of 100 IC_50_. This ensures that all newly produced virions are labeled. The IC_50_ values of most of the Nabs used are less than 50 ng/ml. Hence 100 IC_50_ corresponds to an antibody concentration of at most 5μg/ml or 33 nM. The k_on_ for antibody binding is ~10^5^ M^-1^ s^-1^ (15, 16) while the rate constant for maleimide labeling at an exposed Cys residue is ~10^3^ M-^1^ s-^1^ (17). Hence the ratio of the rates for Env labeling, relative to antibody binding is expected to be (10^3^x10^-2^)/(10^5^×33×10^-9^) i.e. ~3000. As a result, labeling is much faster than binding. Additionally, unlike binding, labeling is an irreversible reaction. Hence, whenever any bound antibody dissociates from Env, labeling at the Cys residue will prevent rebinding. Therefore, the ExCysLib is always expected to be labeled, even in the presence of 100 IC_50_ of the NAb, and there should be virtually no unlabeled virus. In the multi-cycle infectivity assay, the growth kinetics of the labeled ExCysLib (inferred from p24 levels in culture supernatants) was found to be similar in the absence and presence of the bNAb VRC01.This results in an infectivity ratio of ~1, following multiple rounds of infection to select for resistant viruses (Figure 2A). For the other bNAbs too, the infectivity ratio was similarly found to be ~1 (Figure 2B). This conclusively demonstrated that a subset of mutant viruses in ExCysLib became resistant to neutralization upon labeling and suggested that epitopes for these bNAbs are present in the ExCysLib and can be mapped.

**Figure 2.**
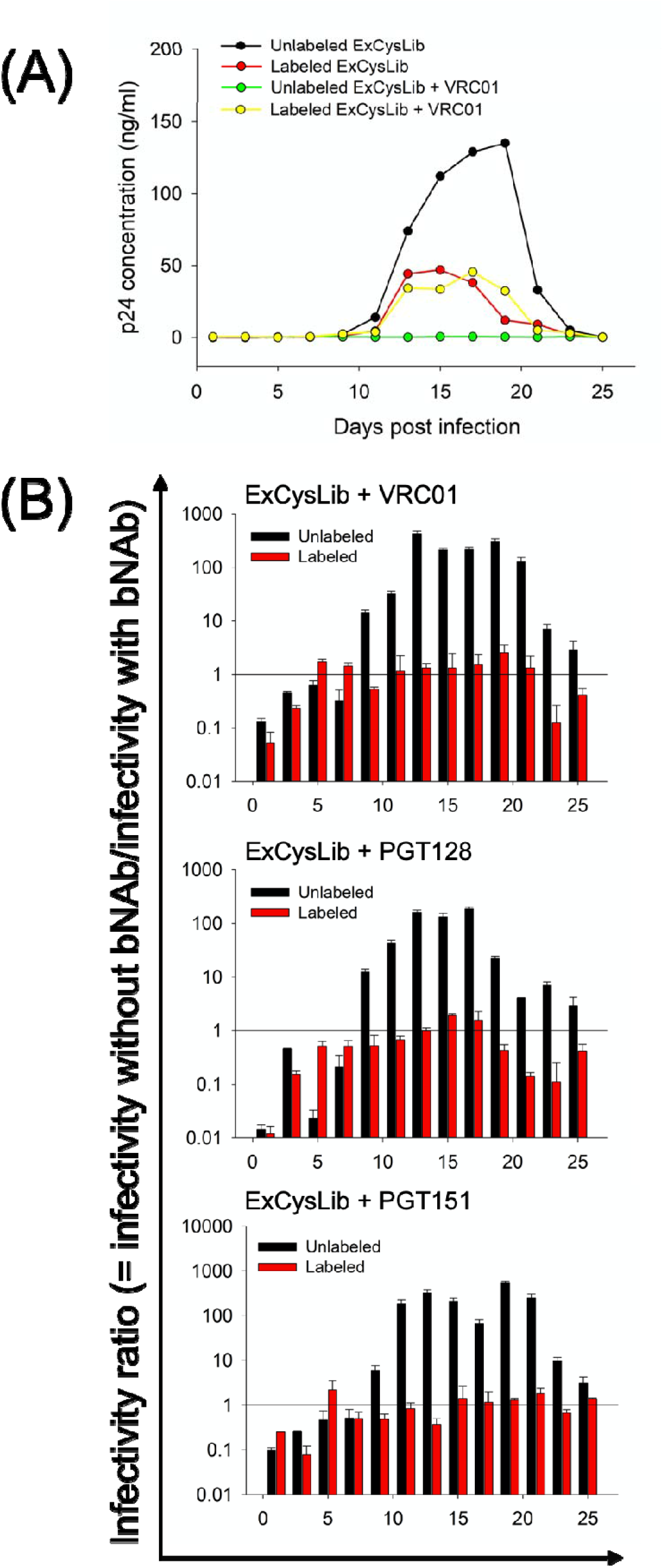
Multi-cycle infectivity of the ExCysLib with bNAbs. **(A) Multi-cycle infectivity of labeled ExCysLib with bNAb VRC01.** The multicycle infectivity of the unlabeled Exposed Cys viral library (black) diminished in the presence of 100 IC_50_ equivalents of bNAb VRC01 (green). Upon labeling of the ExCysLib (red), after multiple rounds of infection in the presence of the labeling reagent, viruses resistant to the bNAb VRC01 emerge (yellow). **(B) Infectivity ratio of labeled ExCysLib with bNAbs.** p24 levels were estimated from multi-cycle infectivity assays of the ExCysLib in the absence and presence of the following bNAbs: VRC01, PGT128 and PGT151, before and after Cys labeling. Here, p24 levels serve as a direct measure of viral infectivity. The infectivity ratio (= infectivity without bNAb/infectivity with bNAb) was straightforwardly used to quantitate viral infectivity changes after Cys-labeling. A ratio of 1 is indicated in the plots as a black line. After ~10 days of infection in the presence of the Cys labeling reagent, the infectivity ratio for labeled ExCysLib with bNAbs is ~1 (red bar), confirming that labeling at exposed, epitope residues on Env prevents neutralization by the corresponding bNAb. In contrast, in the absence of labeling, viral entry is abolished in the presence of the bNAb (black bar), leading to a high value of the ratio (>>1). Infectivity ratios <1 at initial time points are attributed to very low levels of virus and are likely artifactual. Error bars represent SDs of two replicate experiments.

### Polyclonal epitope specificity mapping using ExCysLib

Delineating anti-HIV polyclonal antibody specificities using standard epitope mapping techniques is challenging since the polyclonal antibody repertoire is most often comprised of bNAbs that target multiple epitopes on Env (18, 19). Characterization of epitope specificities of polyclonal antibodies from HIV-1 infected elite neutralizers or vaccinated individuals (20) can aid in the design of better immunogens against the virus (21). Computational algorithms, such as next-generation neutralization fingerprinting (22) have been employed to map polyclonal antibody specificities from simulated sera, but these methods do not yield conclusive results. Other low-throughput epitope mapping techniques such as Pepscan ELISA (23), use of peptides mimotopes (24) and epitope masking (25) also fail to yield accurate epitope information when multiple specificities are targeted by sera.

We therefore extended our methodology to characterize epitope specificities of polyclonal sera elicited by HIV-1 infected individuals and immunized animals. Plasma from a subtype-C HIV-infected elite neutralizer (numeric identity: NAB059) was used for epitope mapping studies using our approach. The NAB059 elite neutralizer exhibited extraordinarily high titers against a panel of Tier-3 pseudoviruses (26, 27), which are the hardest to neutralize. Epitope mapping was also carried out using sera from guinea pigs immunized with an HIV-1 candidate vaccine, a trimeric cyclic permutant (CycP) of gp120 (28) which elicited Tier-2 neutralization activity against a global panel of HIV-1 isolates (29).

Multi-cycle infectivity assays of the labeled ExCysLib were carried out in HUT-R5 cells, in the presence of 100 ID_50_ equivalents of NAB059 plasma or 10 ID_50_ equivalents of trimeric CycP-immunized guinea pig sera. Both the NAB059 plasma and CycP sera were found to select for neutralization resistant viruses in the labeled ExCysLib (Figure S6), indicating that epitopes for these antibodies present in these polyclonal sera are present in our library.

### Deep sequencing of env gene accurately maps NAb epitopes at single residue resolution

Deep sequencing of the env gene recovered from labeled virion pools was carried out, in the absence and presence of NAbs (bNAb/plasma/sera), to map epitopes at single residue resolution. The env gene was sequenced in 6 fragments with PCR primers containing unique MID tags representing a particular selection condition: bNAb (MID 1-3), plasma/sera (MID 4-5) or controls (MID 6-8) (Table S3). A total of 69 Cys mutants were identified from the deep sequencing data. Of the 12 missing mutants out of the original 81 mutants, 10 mutants (D78C, R151C, N160C, N187C, E267C, E293C, D340C, N392C, N398C, R440C) were not present in the plasmid DNA pool (no reads for MID 8) used for transfection and were likely lost while pooling. This could either be because of loss during pooling or due to degradation of a few plasmids upon storage. In future implementations this problem can be avoided by not isolating individual mutants and subjecting to Sanger sequencing, but instead creating the library as a pool (30). The presence of unmutated WT in the pool will not cause any problems as WT virus is neutralization sensitive and will not infect in the presence of neutralizing antibody.

A couple of mutants (N88C and Q442C) were present in the mutant plasmid DNA pool (MID 8) but showed complete loss of infectivity upon Cys labeling (no reads for MID 6). The number of reads for each Cys mutant in the presence of each NAb was normalized with respect to the corresponding number in the absence of NAb, to calculate normalized read ratios (R), as described in the Methods section. R represents the fold enrichment of Cys mutant reads in the presence of NAbs (MIDs 1-5) compared to reads in the absence of NAbs (MID 6). R values were represented using heatmaps (Figure 3B). For virtually all mutants outside the epitope(s), R is close to zero, because antibody-mediated neutralization eliminates these non-epitope Cys mutant viruses. Residues with R > 0.5 were designated as epitope residues for NAbs.Epitopes of bNAbs identified using our methodology (Figure 3B, Table S4) were compared to structurally defined epitopes calculated from Env:bNAb complexes obtained from PDB (Figure 3A). Our method identifies most of the structurally defined epitopes: 100% for VRC01 (7 out of 7), 80% (4 out of 5) for PGT128 and 60% (3 out of 5) for PGT151 (Figure 3, Table S4). There is one false negative for PGT128 (N135), this residue contributes only a small interfacial area of 21 Å^2^. There are two false negatives for PGT151 (V518 and S640), which are structurally defined epitope residues that our methodology does not identify (Table S4). Of these two, V518 has a small interfacial area of 25 Å^2^ while, S640 buries about 35 Å^2^ in the Env:PGT151 complex. In addition, there are two putative false positives for PGT151 (T63 and R633) which are not part of Env:bNAb contacts yet are identified using our methodology (Table S4). Residue 63 in gp120 is apparently disordered in the complex with PGT151, however the present data suggest that it makes contact with the antibody. R633 is located 9.2 Å from the nearest structurally identified epitope residue N637. It is possible that the label at residue 633 perturbs the glycan attached to residue N637. Despite these discrepancies, epitopes identified by us were in good overall agreement with available structural information (Figure 3).

**Figure 3.**
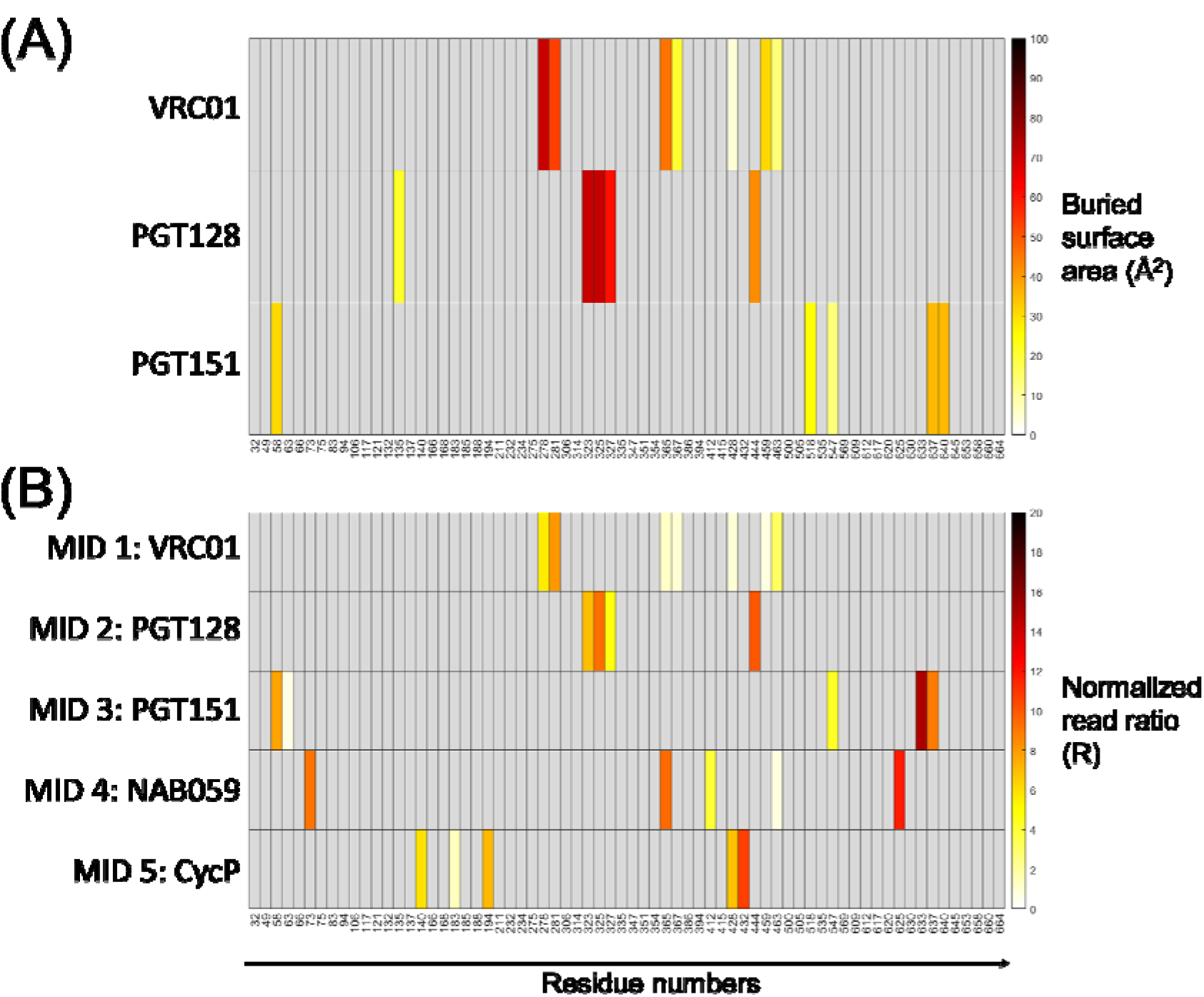
Accurate identification of bNAb and sera/plasma epitopes on Env at single residue resolution. (A) Heatmap showing change in solvent accessible surface area upon binding to selected bNAbs. Accessible surface area for residue side-chains was calculated from crystal structures of complexes of Env with three bNAbs directed to distinct epitopes: VRC01, PGT128 and PGT151. Solvent accessible surface area for residue side-chains was calculated using NACCESS V 2.1.1. The buried surface area was obtained by subtracting the accessible surface area of Env residue side-chains in the presence of the bNAb from the area in the absence of the bNAb. The following PDB IDs were used for surface area calculations: 3NGB (for VRC01), 5C7K (for PGT128) and 5FUU (for PGT151). (B) Heatmap showing normalized read ratios (R) following deep sequencing. Read numbers for selected Cys mutants for the different bNAb selection conditions (MIDs 1-3, corresponding to bNAbs VRC01, PGT128 and PGT151) were obtained from deep sequencing data and normalized by the reads in the unselected condition, in the absence of bNAbs (MID 6), as described in the Methods section. Values > 0.5 show significant enrichment of Cys reads at selected positions, and correspond to predicted epitope residues for the particular bNAbs. Epitopes identified from deep sequencing reads agree well with known epitopes of these bNAbs, obtained from structural information. This data served to validate our strategy. MIDs 4 and 5 correspond to normalized read ratios (R) following deep sequencing for polyclonal plasma and sera respectively. Deep sequencing reads were normalized similar to that for bNAbs. This was directly used to identify unknown epitopes for the NAB059 elite neutralizer plasma (MID 4) and CycP-immunized guinea pig sera (MID 5). All heatmaps show the 69 residues identified from deep sequencing data analysis. Heatmaps were created using MATLAB version 2020a.

Epitopes in plasma from the NAB059 elite neutralizer contained quaternary epitope-specific and CD4-binding site-specific signatures (Figure 3B, Table S4). The CycP sera contained epitope specificities mapping to the V1V2 loop and CD4-binding site (Figure 3B, Table S4), which were consistent with earlier observations using mammalian cell surface display and mutant pseudoviruses (31, 32). Both polyclonal sera were observed to encompass multiple antibody specificities (Figures 3B), which account for their observed neutralization activity.

### Epitope mapping strategy validated using individual Env epitope mutant viruses

Finally, we validated our epitope mapping strategy on a panel of Env-epitope mutants derived from our deep sequencing data (Table S5). Individual viruses containing a Cys mutation either at or outside a bNAb epitope in Env were labeled, and single-cycle viral infectivity assays were performed in TZM-bl cells in the absence and presence of bNAbs. Viral titers were measured using both qRT-PCR (Figure S7, Table S6) and TZM-bl cell-based reporter assays (Figure S8).

Viral infectivity assays of individual Cys mutant viruses were carried out in the presence of 100 IC_50_ equivalents of bNAbs VRC01, b12, PGT128 and PGT151. Viruses became resistant to neutralization only when the corresponding epitopes were masked by the bulky Cys label (Figure 4). Control experiments were carried out by labeling Cys mutant viruses in the presence of bNAbs targeting other sites on Env (Figure 5). These validation experiments clearly demonstrated that our methodology could distinguish between epitope and non-epitope residues for each bNAb.

**Figure 4.**
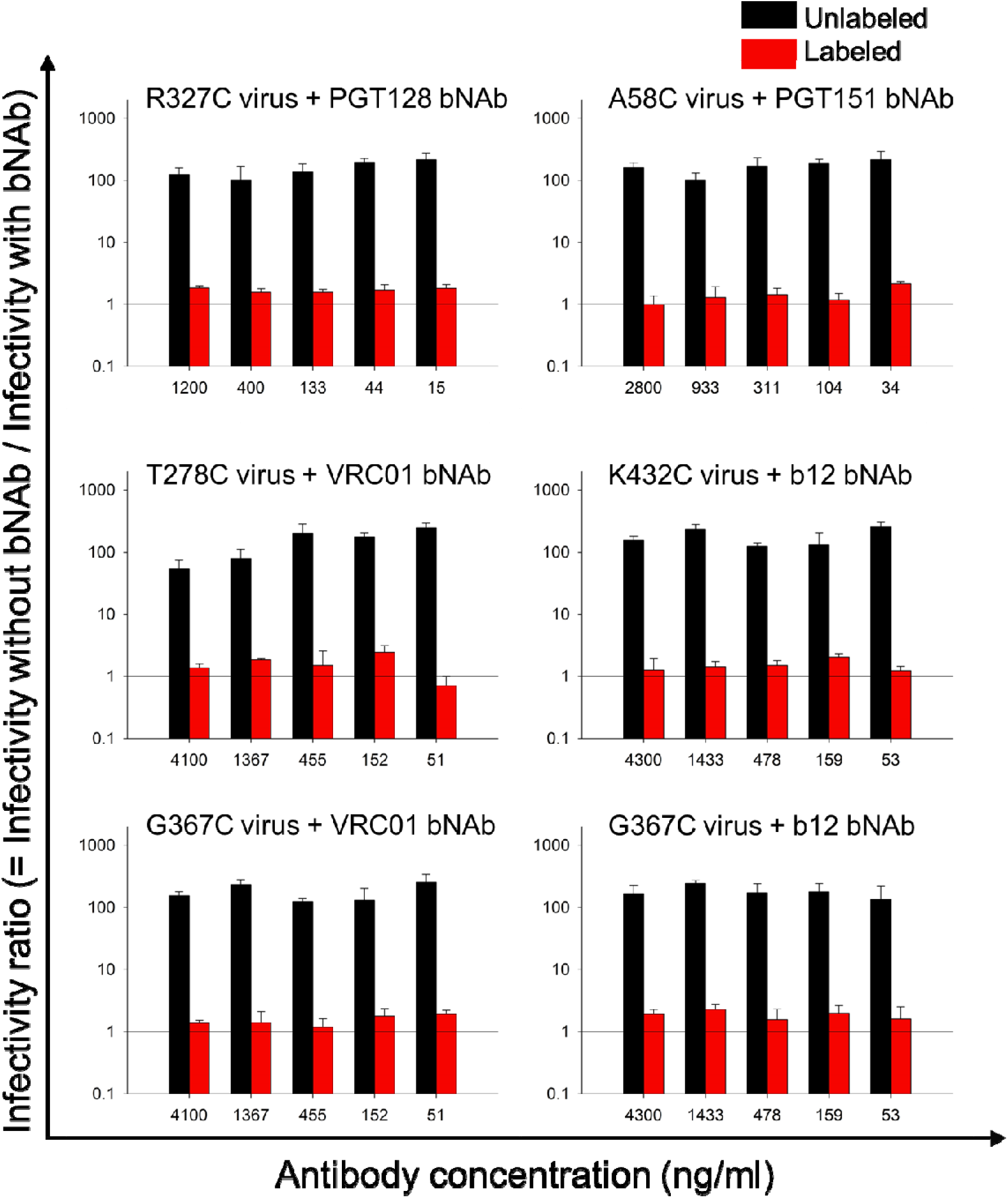
Effect on viral infectivity when the labeled Cys is at the bNAb epitope. If an Env-Cys mutant is part of a particular bNAb epitope, labeling of this residue is expected to block the binding and neutralization of labeled viruses with this specific-Cys mutation. Upon Cys labeling, the Env epitope is masked by the label, and consequently, labeled viruses with Cys mutants at Env-epitope residues become resistant to neutralization by bNAbs targeting that specific epitope. By monitoring the infectivity of labeled and unlabeled Cys mutant viruses in the absence and presence of multiple bNAbs, we were able to confirm epitope targeting by these bNAbs. This was experimentally demonstrated for the following Env-Cys mutant viruses and their corresponding epitope-targeting bNAbs: R327C mutant virus + PGT128, A58C mutant virus + PGT151, T278C mutant virus + VRC01, K432C mutant virus + b12, G367C mutant virus + VRC01, G367C mutant virus + b12. Infectivity (measured as RLUs) was quantified using single-cycle infectivity assays in TZM-bl cells. In the absence of labeling, infectivity ratio (=infectivity without bNAb/infectivity with bNAb) is ~100 fold higher in the absence of antibody (black bars). However, following labeling, viral infectivity is unaffected by the antibody (red bars). Error bars represent SDs of two replicate experiments.

**Figure 5.**
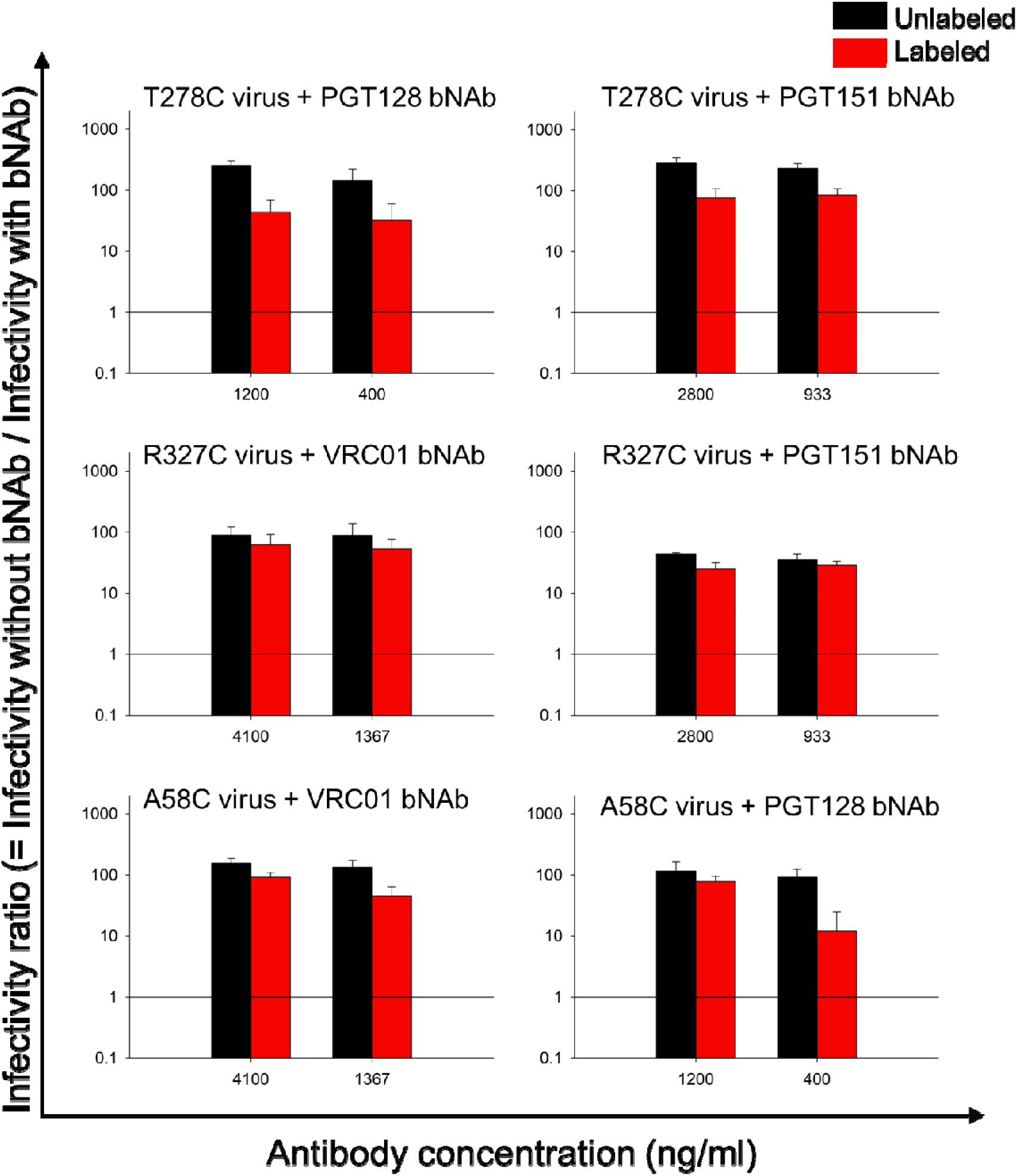
Effect on viral infectivity when the labeled Cys is not at the bNAb epitope. Infectivity of a CD4bs-directed VRC01 epitope mutant virus (T278C) was measured in the presence of other non-CD4bs bNAbs PGT128 and PGT151. Similarly, infectivity of a V3 loop-directed PGT128 epitope mutant virus (R327C) was measured in the presence of bNAbs with different, non-overlapping epitopes: VRC01 and PGT151. Further, infectivity of a gp120:gp41 interface directed PGT151 epitope mutant virus (A58C) was monitored in the presence of bNAbs targeting different epitopes: VRC01 and PGT128 bNAbs. In all these cases, the Cys-Env mutant is not the epitope of these bNAbs, and both before and after labeling, mutant viruses were found to be similarly sensitive to neutralization. This was inferred from the infectivity ratio of the labeled viruses (= infectivity in the absence of bNAb/ infectivity in the presence of bNAb) which was >> 1. Infectivity (measured as RLUs) was quantified using single-cycle infectivity assays in TZM-bl cells. Error bars represent SDs of two replicate experiments.

## Discussion

High-throughput, site-directed scanning mutagenesis techniques are widely used in protein engineering. These enable studies of protein structure and function at single residue resolution (1, 2, 33, 34). A deep mutational scanning-based approach, “mutational antigenic profiling” was recently used to map epitopes of nine HIV-1 bNAbs (35). This approach is based on the creation of codon-mutant libraries (36), identification of bNAb escape mutants and deep sequencing of virus pools to quantify the enrichment of mutations after antibody selection. This method comprehensively delineates regions of viral escape or functional epitopes, not all of which are in physical contact with the antibody (35).

We used site-directed Cys scanning mutagenesis, a powerful tool to probe protein function (37–40). Cys is an attractive candidate for mutagenesis since Cys at solvent-exposed sites in proteins can be labeled site-specifically. In addition, Cys is relatively infrequent in the proteome, and introduction of Cys in proteins is most often accompanied by very little structural and functional perturbation (41–43). Moreover, our method is devoid of the complexity of saturation mutagenesis libraries, which makes library construction and analysis of deep sequencing data considerably simpler. Our library sizes are at least 20-60 times smaller than saturation mutagenesis libraries. Our method appears to exclusively identify structurally defined epitope residues that are in close physical proximity to the antibody (Figure 3). Recently developed cryo-EM based epitope-mapping approaches (44) are also useful for mapping monoclonal antibody epitopes. However, for polyclonal sera, the cryo-EM approach identifies the predominant epitopes targeted. In the absence of any enrichment strategy, these are unlikely to be neutralizing epitopes, since neutralizing antibodies typically make up a small fraction of the total antibody response (45). The present methodology is similar in overall principle, to recently developed saturation mutagenesis approaches (13, 35, 36) to identify neutralization epitopes. The present approach, when combined with our earlier cell surface labeling approach which elucidates all the epitopes targeted in the response (1), provides a comprehensive snapshot of the regions of the immunogen surface targeted by antibodies as well as the fraction of the surface targeted by neutralizing antibodies. The present method does not require sophisticated instrumentation, is highly parallelizable, can map epitopes to single residue resolution and throughput can be further improved by using barcoded libraries. These features make it an attractive approach to map neutralization responses in human sera from either HIV-1 infected individuals or vaccinees. Knowledge of the three-dimensional structure of the antigen is not a prerequisite for selecting residues for mutagenesis using the proposed methodology. In the absence of structural information, homology modeling or Asp scanning mutagenesis (46, 47) can be employed to locate solvent accessible residues in proteins. The methodology can readily be coupled to cell surface display methodologies (8) to obtain all, rather than just neutralization epitopes. A limitation of the present work is that any Cys mutants at positions where labeling results in loss of viral infectivity will be lost from the library and cannot be studied. We did observe a couple of mutations (N88C, Q442C) that were present in the plasmid DNA pool but showed loss of infectivity upon labeling. Cys mutants at these residues likely interfere with viral functions such as CD4-binding or viral fusion and were therefore eliminated in our methodology. The Cys label, Maleimide-PEG2-Biotin, has a long arm length of 29.1Å. While the bulk of the label blocks bNAbs from binding, it can potentially lead to the identification of a few false positives, this needs to be confirmed through additional experiments. Finally, in this initial implementation, a subset of surface exposed residues was mutated to Cys, to minimize the size of the library while maintaining relatively uniform surface coverage. In future implementations, all surface residues can be mutated, which would increase the library size, but would result in an enhanced resolution of the inferred epitopes.

There is an urgent need for reliable epitope mapping techniques to elucidate neutralizing epitopes against HIV-1 and others emerging viral infections, including SARS-CoV-2. Deep sequencing of the env gene recovered from labeled virion pools in the presence of bNAbs/sera, using our methodology, provides comprehensive knowledge of epitope specificities, which in turn provides useful insights about the nature of antibody response elicited by the virus (48–50) and informs immunogen design (51, 52).

## Materials and Methods

### Plasmids

Site-directed Cys scanning mutagenesis was carried out in the env gene (~2.2 kb) of HIV-1 molecular clone pLAI-JRFL (11.7 kb), where LAI-JRFL refers to the strain of HIV-1. Since site-directed mutagenesis is less efficient for larger plasmids (>3.1 kb), the env gene was subcloned in a smaller TA cloning vector pTZ57/R.

### bNAbs

All bNAbs were obtained from the IAVI Neutralizing Antibody Consortium or the NIH AIDS reagent program (https://www.aidsreagent.org/).

### Residue selection for ExCysLib

A combined criterion of >30% solvent accessibility and sidechain-sidechain centroid distance >8 Å was used to select 81 residues from the X-ray crystal structure of the pre-fusion HIV-1 Env ectodomain (PDB ID: 4TVP). The solvent accessibility of each residue was calculated using the NACCESS V 2.1.1 program (http://wolf.bms.umist.ac.uk/naccess/) after removing antibody chains. Pairwise sidechain centroid distances were calculated for all residues exhibiting solvent accessibility >30% using an in-house PERL script. This side-chain side-chain centroid distance cut-off was chosen to leave out residues in close proximity (8). Selected residues occupy a surface area roughly corresponding to one-third of the total surface area of the Env ectodomain (PDBePISA, EMBL-EBI). In addition, 47 out of the 81 residues are contact sites either for CD4 or the following bNAbs directed to various neutralizing epitopes on Env: CD4-binding site (VRC01, PGV04, 3BNC117, NIH45-46, CH103, b12), CD4-induced epitope (17b), V3-glycan (PGT121-123, 125-128, 130, 131, 135), V1V2-glycan (PG9, CAP256-VRC26), MPER (2F5), gp120-gp41 interface (PGT151) and mannose-dependent epitope (2G12).

### Generation of Exposed Cys mutant plasmid library

81 selected exposed residues (Table S1) were individually mutated to Cys by site-directed mutagenesis using partially overlapping complementary primers which generates a doubly-nicked amplicon (30). There was a 21 bp overlap between the forward and reverse primers with a Cys codon 9 bp from the 5’-end of each primer. Primers were synthesized at the Protein and Nucleic Acid facility at Stanford University, USA. Mutagenesis was carried out using Phusion DNA polymerase (NEB) according to the manufacturer’s protocol in a thermal cycler PTC-200 (Bio-Rad). Each 20 μl reaction had 20-25 ng of template pTZ57/R-env, 1 μM of the primer pair, 1X GC Buffer and 200 μM dNTPs, with 3% DMSO as an additive. PCR was done for 30 cycles at an annealing temperature of 65°C for 35 seconds. 8.5 μl of the PCR product was digested with 0.5 μl DpnI (20 U/μl) and 1 μl CutSmart buffer at 37°C for 6 hours. PCR products were purified using the GeneJET PCR Purification kit (Thermo Fisher Scientific). 1 μl of the purified PCR product was used to transform *E.coli* TOP10 electrocompetent cells. Plasmids were isolated using commercially available miniprep columns (Qiagen). Presence of the Cys mutation at selected positions in env was confirmed by Sanger sequencing (at Macrogen, South Korea).

Single-Cys mutant pTZ57/R-env plasmids were pooled in equimolar quantities. The mutant env gene was amplified from pTZ57/R-env plasmid pools using Phusion DNA Polymerase (NEB) and primers from 5’ and 3’ ends. This was cloned into the pLAI-JRFL molecular clone between BamHI and SfiI restriction sites using T4 DNA ligase (NEB), thus replacing WT env. For this, the vector backbone fragment was purified by electroelution (at 100 volts in 0.5X TAE for 1.5 hours), extracted using phenol-chloroform-isoamyl alcohol (25:24:1), precipitated using Pellet Paint Co-Precipitant (EMD Millipore), and finally, ligated and electroporated into *E. coli* TOP10 cells. The Exposed Cys mutant plasmid library was purified using commercially available miniprep columns (Qiagen) using growth conditions of 30°C and 130 rpm to support the propagation of large plasmids.

### Transient production of ExCysLib

The ExCysLib was produced by transient transfection of the Exposed Cys mutant plasmid library into HEK293T cells. 24 hours before transfection, 5×10^5^ cells/well were seeded in 6-well plates containing 2 ml of DMEM with 10% Fetal Bovine Serum (10% DMEM). The transfection mix contained 16 μg of transfection reagent polyethyleneimine (PEI) (Sigma-Aldrich) and 4 μg of mutant pLAI-JRFL plasmid (4:1 ratio of PEI:DNA), in a final volume of 100 μl of serum-free DMEM. After 10 minutes of incubation at room temperature, 100 μl of 10% DMEM was added to the transfection mix and used to transfect HEK293T cells at 50-70% confluency. 6-8 hours post-transfection, media was replaced with 1.5-1.8 ml of 10% DMEM, and 48 hours post-transfection, culture media was collected to harvest the ExCysLib. Culture media was centrifuged at 1500 rpm for 10 minutes to remove residual cells. To the viral supernatants, FBS was added at a final concentration of 20%. ExCysLib was aliquoted and stored at −70°C. All studies with replication competent viruses were carried out at the Biosafety Level-3 facility at the Centre for Infectious Disease Research (CIDR) at IISc.

### Single-cycle viral infectivity of ExCysLib in TZM-bl cells

Single-cycle infectivity of ExCysLib was measured using a luciferase reporter-based assay in TZM-bl cells (53). 100μl of undiluted and threefold-diluted ExCysLib in 10% DMEM, was added in successive wells of a 96-well tissue culture plate. To this, 1×10^4^ freshly-trypsinized TZM-bl cells (in 100 μl of 10% DMEM containing 30 μg/ml of DEAE-dextran) was added. 48 hours post-infection, 100 μl volume was discarded from each well and 80μl of Britelite Plus reagent (PerkinElmer) was added and incubated for 2 minutes at room temperature. The reagent carries out cell lysis as well as the luciferase enzymatic reaction. Following incubation, 130 μl was aspirated from each well, transferred to a 96-well black plate and luminescence, measured in relative luminescence units (RLUs), was recorded in a VICTOR X2 luminometer (PerkinElmer).

### Quantification of Cys labeling using MALDI-TOF

WT CcdB and the test protein, GyrA14-I491C were purified using previously standardized protocols (54). GyrA14-I491C was dissolved at 50 μM in 1X PBS buffer containing 20% FBS (to mimic virus labeling) at pH 7.0-7.5, treated with 0.5mM TCEP to reduce disulfide bonds, and labeled with a commercially available Cys label, EZ-Link™ Maleimide-PEG2-Biotin (Thermo Fisher Scientific). At pH 7.0, the maleimide group is almost 1000 times more reactive towards free sulfhydryl groups than to primary amines in the protein (55). Labeling was carried out for different concentrations of the label, i.e. 0.01 mM, 0.1 mM, 1 mM and 10 mM and different time-points of incubation i.e., 15 mins, 1 hour, 4 hours and overnight; in a final labeling volume of 100 μl at 4°C. Higher concentrations of the label exceeded maximum solubility, and hence could not be used. Following labeling, excess label was removed using a PD Minitrap G-25 desalting column (GE Healthcare). The labeled GyrA14-I491C was loaded onto a pre-equilibrated CcdB affinity column prepared using Affi-gel 10 (Bio-Rad), and incubated at 4°C for 4 hours on a rotary mixer to enable binding. Unbound protein (containing components of FBS) was removed by washing the column with 10 times the bed volume of coupling buffer (0.05M sodium bicarbonate, 0.5 M NaCl, pH 8.3), and eluted using 0.2 M Glycine, pH 2.5 into a vial containing an equal volume of 400 mM HEPES, pH 8.5. Mass spectra of the elutes were acquired on an UltrafleXtreme MALDI TOF (Bruker Daltonics). Mass of unlabeled test protein was 17.5 kDa, and upon labeling, an increase in mass of ~0.5 kDa was observed, corresponding to the mass of the Cys label. The approximate extent of Cys labeling of the GyrA14-I491C test protein was calculated as:

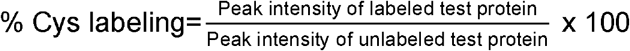

### Cys labeling of WT virus

Transient transfection of WT pLAI-JRFL plasmid into HEK293T cells led to the production of replication competent WT LAI-JRFL virus. All virus stocks were made cell-free by filtration through a 0.45 μm syringe filter. 3×10^5^ RLU equivalents (as determined by TZM-bl titrations) of WT virus was labeled with 1mM and 10 mM EZ-Link™ Maleimide-PEG2-Biotin in 1X PBS, overnight, in a reaction volume of 100 μl at 4°C. Post-labeling, single-cycle viral infectivity was measured in TZM-bl cells.

### Neutralization potency (IC_50_) of WT virus with bNAbs

The neutralization potency (IC_50_) of the WT LAI-JRFL virus with bNAbs VRC01, b12, PGT128 and PGT151 was measured using viral neutralization assays in TZM-bl cells. This assay measures the antibody-mediated neutralization of HIV-1, by quantifying the reduction in RLUs i.e. Tat-driven luciferase gene expression upon viral infection. 1-1.5×10^5^ RLU equivalents of WT LAI-JRFL virus were incubated with serial threefold-dilutions of bNAbs for 1 hour at 37°C in a total volume of 100 μl of 10% DMEM. To this, 1×10^4^ freshly-trypsinized TZM-bl cells in 100μl containing 30μg/ml of DEAE-dextran were added, and incubated for 48 hours. Luminescence was measured using Britelite Plus (PerkinElmer) on a VICTOR X2 luminometer (PerkinElmer). Neutralization potency (IC_50_) is calculated as the sample dilution at which the RLUs in test wells (with bNAb) are reduced by 50% in comparison to RLUs in virus control wells (without bNAb) after subtraction of background RLUs from cell control wells. We used concentrations 100 times in excess of the calculated IC_50_ with different bNAbs (12, 13) for subsequent epitope-mapping experiments.

### Multi-cycle infectivity of ExCysLib

We carried out multi-cycle infectivity assays of the ExCysLib in HUT-R5 cells (a derivative of the human T-cell line HUT-78, stably transduced with the CCR5 coreceptor). This assay evaluates both infectivity as well as the rate and efficiency of virus production in cell culture over a course of time (replication fitness). 5×10^6^ HUT-R5 cells were infected with 3×10^5^ RLU equivalents (estimated from TZM-bl titrations) of ExCysLib using 30 μg/ml DEAE-dextran in a total volume of 250 μl of 10% RPMI 1640. Infection was carried out by incubation in a 37°C CO_2_ incubator for 2 hours, with occasional shaking every half-hour. Excess virus from infected cells was removed by washing with 45 ml of ice-cold 1X PBS. ExCysLib-infected cells were centrifuged at 1200 rpm for 5 mins, the supernatant was discarded, the cell pellet was resuspended in 6 ml of 10% RPMI 1640, transferred to a T-25 flask and kept in a CO_2_ incubator. Cells were split in a 1:2 ratio every alternate day post-infection, and virus replication was quantitated by measuring the p24 antigen released in culture supernatants at different time points using HIV-1 p24 SimpleStep ELISA kit^®^ (Abcam). Serial dilutions of HIV-1 p24 protein (strain 92418, subtype B) provided with the kit was used to plot a standard curve. Multi-cycle infectivity of WT virus and ExCysLib was measured by estimating p24 levels from the standard curve.

### Multi-cycle infectivity of labeled ExCysLib in the presence of monoclonal bNAbs

To map epitopes of known bNAbs using our methodology, we carried out multi-cycle infectivity assays of the labeled ExCysLib in the presence of 100 IC_50_ equivalents of bNAbs VRC01, PGT128 and PGT151. This was carried out by labeling the ExCysLib, with 10mM Maleimide-PEG2-Biotin for 2 hours at 37°C, followed by incubation with bNAbs for 1 hour at 37°C. This was followed by multi-cycle infectivity assays in HUT-R5 cells. For this, every alternate day post-infection, cells were split 1:2 with RPMI growth media and the corresponding culture supernatants containing Cys mutant viruses (ExCysLib) were labeled with fresh 10mM Maleimide-PEG2-Biotin for 2 hours. Next, sufficient bNAb was introduced into the culture media to keep final bNAb titers at a value of 100 IC_50_. Since epitopes for selected bNAbs are present in the ExCysLib, some mutant viruses are expected to be resistant to neutralization upon labeling and subsequent bNAb exposure. This was inferred after multiple rounds of infection to enrich such mutants both by quantifying the ratio of p24 levels of the labeled ExCysLib in the absence to presence of bNAb and by deep sequencing.

### Neutralization titer (ID_50_) of polyclonal human plasma and guinea pig sera

The neutralization titer or 50% inhibitory dose (ID_50_) was determined using TZM-bl cell-based viral neutralization assays. Neutralization assays was carried out for the WT LAI-JRFL virus in the presence of plasma from the elite neutralizer, NAB059 and for terminal bleed sera (collected at week 34.5) from guinea pigs immunized with CycP gp120 trimers. Similar to IC_50_, neutralization titer, or 50% inhibitory dose (ID_50_) is calculated as the reciprocal of the serum dilution at which the RLUs in test wells (with plasma/sera) are reduced by 50% in comparison to RLUs in virus control wells (without plasma/sera) after subtraction of background RLUs from cell control wells. Titers 100 times in excess of the calculated ID_50_ were used to map epitopes of NAB059 plasma using the ExCysLib, as discussed above for the monoclonal antibodies. For mapping epitopes using the immunized guinea pig sera, 10 times the ID_50_ titer was used, since limited amounts of this sera was available.

### Multi-cycle infectivity of labeled ExCysLib with polyclonal plasma/sera

To map epitopes in polyclonal sera, multi-cycle infectivity assays (56) of the labeled ExCysLib was carried out with 100 ID_50_ equivalents of NAB059 plasma or 10 ID_50_ equivalents of immunized guinea pig sera, in HUT-R5 cells. Multi-cycle infectivity assays of labeled virus in the presence of plasma/sera were performed as described for bNAbs. As with the bNAbs, following each round of cell splitting, Maleimide-PEG2-Biotin labeling reagent was added to a final concentration of 10mM. After two hours, additional plasma/sera was introduced into the culture media to keep final neutralizing titers constant at 100 IC_50_. Finally, the ratio of p24 levels of the labeled ExCysLib was quantified in the absence to presence of plasma/sera at each time point.

### Deep Sequencing of env gene to map epitopes

Replicative virus isolated from HUT-R5 culture supernatants on day 13 post-infection was used to isolate RNA and prepare cDNA using RT-PCR. Viral RNA was isolated from 500 μl of tissue culture supernatant of HUT-R5 cells using a QIAamp^®^ viral RNA Mini Kit (Qiagen). RNA yield and purity were quantitated using NanoDrop™ 2000. SuperScript™ III reverse transcriptase (Thermo Fisher Scientific) was used to produce cDNA from viral RNA using a primer (pLAI-JRFL-NefOR-rev) which binds 811 bp downstream of the *env* gene. The primer pLAI-JRFL-NefOR-rev has the sequence 5’-AGGCAAGCTTTATTGAGGC-3’. First-strand cDNA synthesis was carried out from the purified viral RNA using 100 ng of viral RNA, 1 μl (200 units) of SuperScript™ III reverse transcriptase, 5 μl of 5X First-Strand Buffer, 0.5 μM of primer pLAI-JRFL-NefOR-rev, 1 μl of dNTP mix (containing 10 mM of each dNTP) and 1 μl RNasin ribonuclease inhibitor (Promega) in a total reaction volume of 25 μl. cDNA synthesis reaction was carried out at 42°C for 50 mins, followed by inactivation of reverse transcriptase at 70°C for 15 mins. The env gene was amplified using Phusion^®^ High-Fidelity DNA Polymerase (NEB) using Phusion HF buffer and primers specific to the 5’ and 3’ ends of *env*, pLAI-JRFL-env-F (5’-TAGGCATCTCCTATGGCAGGAAG-3’) and pLAI-JRFL-env-R (5’-GTCTCGAGATGCTGCTCCCACCCC-3’). PCR was done for 30 cycles with an annealing temperature of 60°C for 35 seconds. Amplified env cDNA was purified using a GeneJET PCR purification kit (Thermo Fisher Scientific). This was further amplified for deep sequencing. The length of the env gene is ~2.2 kb, whereas the maximum length of DNA that can be sequenced on the Illumina HiSeq 2500 platform is 500 bases in a paired-end sequencing run (2×250 bp). Because of this size restriction imposed by Illumina sequencing, the env gene was sequenced in six fragments.

The env gene was sequenced from the ExCysLib isolated under different conditions (listed in Table S3). Primers were designed for each of the conditions, with a unique Multiplex IDentifier (MID) tag representing each condition. For the first 5 fragments both the forward and reverse flanking primers begin with NNN, followed by 6 bases of the unique sequence tag for each MID and 21 bases complementary to the gene. The sixth fragment contained the same flanking forward primer, but the reverse primer was devoid of the MID sequence. Each MID sequence corresponds to one of the following conditions: 3 broadly neutralizing antibodies (bNAbs VRC01, PGT128, PGT151), 2 sera (NAB059 patient sera and CycP guinea pig sera) and 3 controls (conditions 6-8, Table S3). The PCR products have an average read length of 420 bp, with an overlap of ~63 bp between successive fragments. PCR for the fragments was carried out using Phusion polymerase with a template concentration of 10ng/μl for 15 cycles. PCR products were quantified by densitometry analysis using the Bio-Rad Quantity One software (Bio-Rad). Fragments were pooled in an equimolar ratio, gel purified and sequenced on the Illumina HiSeq 2500 platform.

### Deep Sequencing Data Analysis

The deep sequencing data analysis pipeline was built using PERL with certain modifications to an already existing, in-house protocol (https://github.com/skshrutikhare/cys_library_analysis). The raw data consisting of 134,897,599 x 2 paired-end reads of the mutants was assembled using PEAR v0.9.6 (Paired-End Read Merger) tool (57). Out of this, 1.102% reads were discarded, leaving 133,410,630 assembled reads. Initial screening of env reads was based on the unique primer pairs for each fragment and was done separately. The same set of MIDs was used for each fragment. A Phred score cut-off of ≥ 20 was applied on these reads for quality filtering. After quality filtering, reads not having the correct MID/primer set for env data were eliminated, along with reads having mismatched MIDs. Thus, each filtered read had the MID, forward/reverse primer sequence followed by the gene sequence. Reads in each fragment were segregated into different bins based on the MID sequence. Before binning, reads with insertions/deletions, with incomplete primer/MIDs (due to quality filtering) and those having incorrect pairs of primers were removed. Binned reads were aligned with the WT sequence of the env gene corresponding to each fragment using Water v 6.4.0.0 program (58). Aligned reads were reformatted for ease of further processing. Finally, these reads were classified based on whether they were WT, or contained indels and substitutions (single, double, triple and multiple mutants). Single Cys mutants were sorted based on read numbers. Normalization was carried out as follows: for Residue i in Fragment x at MID y, where x = 1 – 6, and y = 1 - 5

Numerator (N) = (# Cys reads (i) / (Total (#Cys reads) + WT reads)) for Fragment x at MID y

Denominator (D) = (# Cys reads (i) / Total (#Cys reads) + WT reads)) for Fragment x at MID 6. Normalized Read Ratio (R) = N/D. All residues with R > 0.5 were designated as epitope residues.

Alternative methods of normalization, for example, using the read ratio of a given Cys mutant at MID x, where x = 1-5 to that at MID 6 yielded very similar conclusions. R represents the relative enrichment of Cys reads in the presence of NAbs, accounting for the WT reads for each MID.

### Viral titer estimation using qRT-PCR of *pol* gene

Viral titer (or viral load) of individual bNAb-epitope Cys mutant viruses was measured using qRT-PCR of the conserved HIV-1 *pol* gene (201 bp) using SYBR Green detection chemistry. To this end, serial dilutions of the pLAI-JRFL plasmid (ranging from 10^1^ to 10^8^ copies/μl) were made in nuclease-free water containing 0.1 μg/μl BSA. qRT-PCR reactions had 10 μl of 2X iQ™ SYBR^®^ Green Supermix (Bio-Rad), 0.5 μM each of *pol* gene-specific forward and reverse primers, 1 μl of the diluted plasmid DNA template and 2 μg of BSA, in a total volume of 20 μl. Sequences of the primers used to amplify the *pol* gene: pLAI-JRFL-pol-forward: 5’-AGGAGCAGAAACGTTCTATGTAGATGG-3’ and pLAI-JRFL-pol-reverse: 5’-GCATATTGTGAGTCTGTTACTATATTTACT-3’. The assay was performed using an iQ5 RT-PCR detection system (Bio-Rad) under the following cycling conditions: initial denaturation at 94°C for 5 mins, followed by 40 cycles of denaturation at 94°C for 30 secs, primer annealing at 60°C for 35 secs, and primer extension at 72°C for 30 secs. The threshold cycle (C_T_) values were obtained from the fluorescence data during amplification and used to plot a standard curve for the different dilutions of pLAI-JRFL plasmid DNA. Melting curve analysis and agarose gel electrophoresis of qPCR products was also carried out to confirm that the observed amplification was specific.

The standard curve was used for the absolute quantification of viral cDNA. To this end, viral RNA was isolated from 500 μl of tissue culture supernatant from HEK293T cells using a QIAamp^®^ viral RNA Mini Kit (Qiagen). RNA yield and purity were quantitated spectrophotometrically using NanoDrop™ 2000. First-strand cDNA synthesis was carried out from the purified viral RNA in a 25 μl reaction containing 100 ng of viral RNA, 1 μl (200 units) of SuperScript™ III reverse transcriptase (Thermo Fisher Scientific), 5 μl of 5X First-Strand Buffer (Thermo Fisher Scientific), 0.5 μM of *pol* gene-specific primer (pLAI-JRFL-pol-reverse), 1 μl of dNTP mix (containing 10 mM of each dNTP) and 1 μl RNasin ribonuclease inhibitor (Promega). The reaction was carried out at 42°C for 50 mins, followed by heat inactivation of reverse transcriptase at 70°C for 15 mins. To estimate viral titers, 2μl of viral cDNA from the first-strand reaction was used for absolute quantification using the standard curve. For this, qRT-PCR of the *pol* gene was carried out using the same reaction components and cycling conditions used earlier for plasmid DNA, except that viral cDNA was used as the template. The C_T_ value obtained from the qRT-PCR data was used to estimate viral copy numbers from the standard curve. An important assumption of this method is that the reverse transcription step is 100% efficient. Although the qRT-PCR assay is capable of accurately measuring viral titers, it does not accurately measure infectious titers, since this assay also accounts for defective particles that harbor non-native forms of the Env and consequently, do not give rise to productive infection.

### Determination of infectious viral titer using TZM-bl cells

Infectious titer of individual bNAb-epitope Cys mutant viruses from HEK293T culture supernatants was estimated using single-cycle infectivity assays in TZM-bl cells. Using the viral titers from qRT-PCR, 3×10^5^ virions were used to infect 5×10^5^ TZM-bl cells, at a resultant multiplicity of infection of 0.6. Finally, single-cycle infectivity was quantified by luminescence measurements from the cells 48 hours post-infection.

## Author Contributions

RD and RV designed the study, interpreted and analyzed data, and wrote the manuscript. RD performed experiments. RRC helped with cloning Cys mutants and performed PCRs of the env gene for deep sequencing. KM processed deep sequencing data. LEH provided reagents which enabled the study.

## Acknowledgements

The authors would like to thank the NIH AIDS reagent program and the IAVI Neutralizing Antibody Consortium for providing HIV antibodies and reagents that enabled the study. Dr. Uddhav Timilsina and Dr. Ritu Gaur from the South Asian University, New Delhi are acknowledged for providing the HUT-R5 cells. The authors are grateful to Jyothi Prabha G and Afroze Chimthanawala for providing purified GyrA14-I491C protein and CcdB protein respectively, and Dr. Shruti Khare for running PERL scripts to calculate pairwise side-chain centroid distances. Dr. Ujjwal Rathore and Dr. Shahbaz Ahmed are acknowledged for helpful discussions. RD and RRC are the recipients of a fellowship from the Council of Scientific and Industrial Research, Government of India. KM is thankful to DST-SERB for financial support. This work was funded by NIH grant no. R01AI118366-01, dt. 07/15/2015 to RV. We also acknowledge funding for infrastructural support from the following programs of the Government of India: DST-FIST, UGC Center for Advanced Study, DBT-IISc Partnership Program, and a JC Bose Fellowship from DST to RV.

## Competing Interests Statement

RD and RV are inventors of a patent on mapping protein binding sites and conformational epitopes using Cys labeling (PCT WO/ 2018/104967).

**Supporting Information** is available.

