## Supplementary Information for "A facile method of mapping HIV-1 neutralizing epitopes using chemically masked cysteines and deep sequencing"

**This PDF file includes:**

Supplementary Text

Figures S1 to S8

Tables S1 to S6

SI References

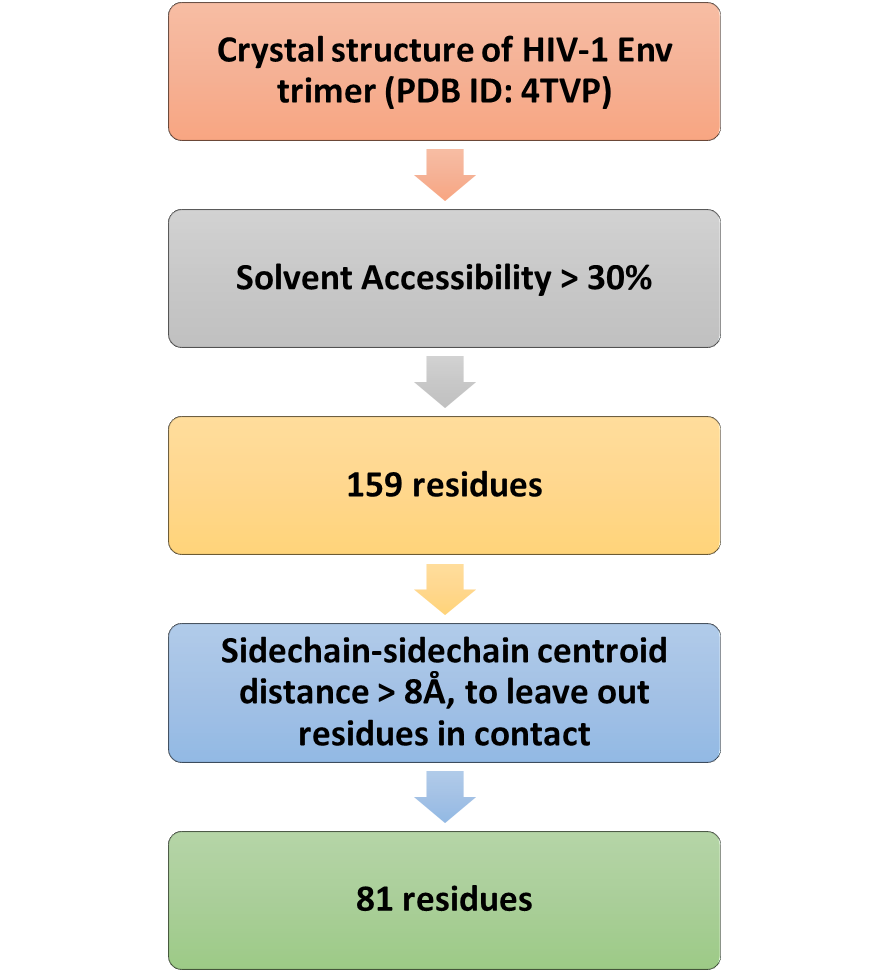

Fig. S1. Residue selection workflow for Exposed Cys Library (ExCysLib).

A total of 81 solvent-exposed residues were selected from the X-ray crystal structure of the prefusion HIV-1 Env ectodomain (PDB ID: 4TVP) using a combined criterion of >30% solvent accessibility and sidechain-sidechain centroid distance >8Å. The solvent accessibility of each residue was calculated using the Naccess V 2.1.1 program. Pairwise centroid distances between sidechains were calculated for all residues exhibiting solvent accessibility >30% using in-house PERL scripts.

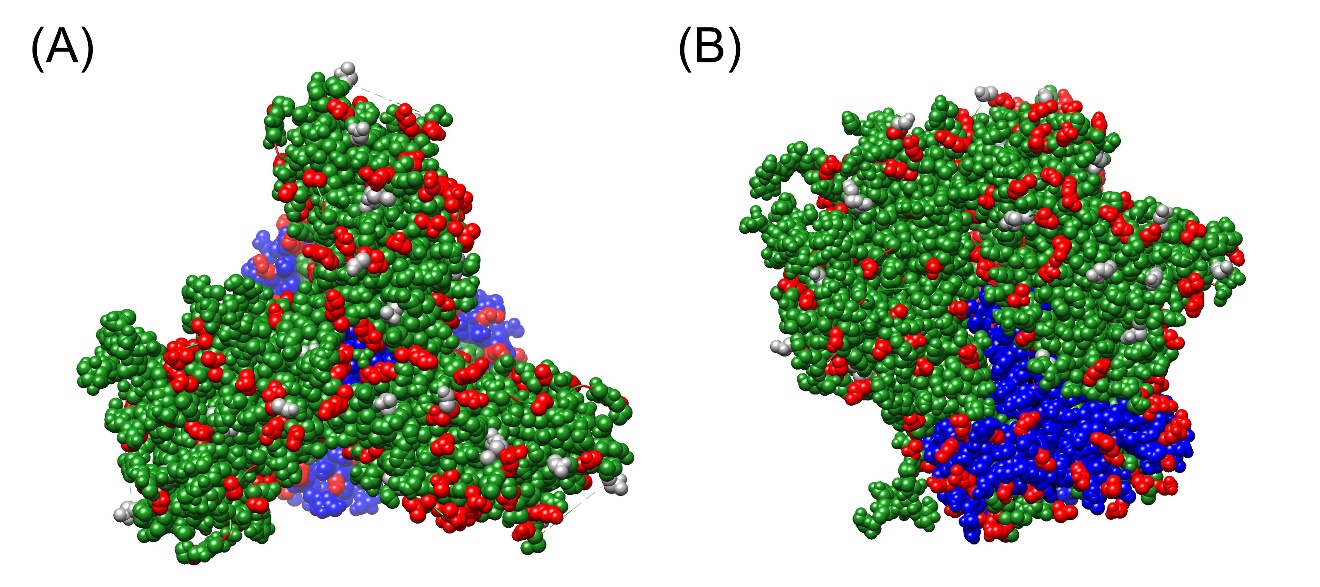

Fig. S2. Selected ExCysLib residues mapped onto the trimeric Env crystal structure (PDB ID: 4TVP) showing (A) top and (B) side view

The three gp120 chains are in green, the three gp41 chains are in blue, the 69 (out of 81 selected) residues identified from the deep sequencing data are shown as red spheres while the rest 12 (out of 81) residues not present in our deep sequencing data are shown as dark gray spheres. Selected residues exhibit substantial and relatively uniform surface coverage of the Env ectodomain.

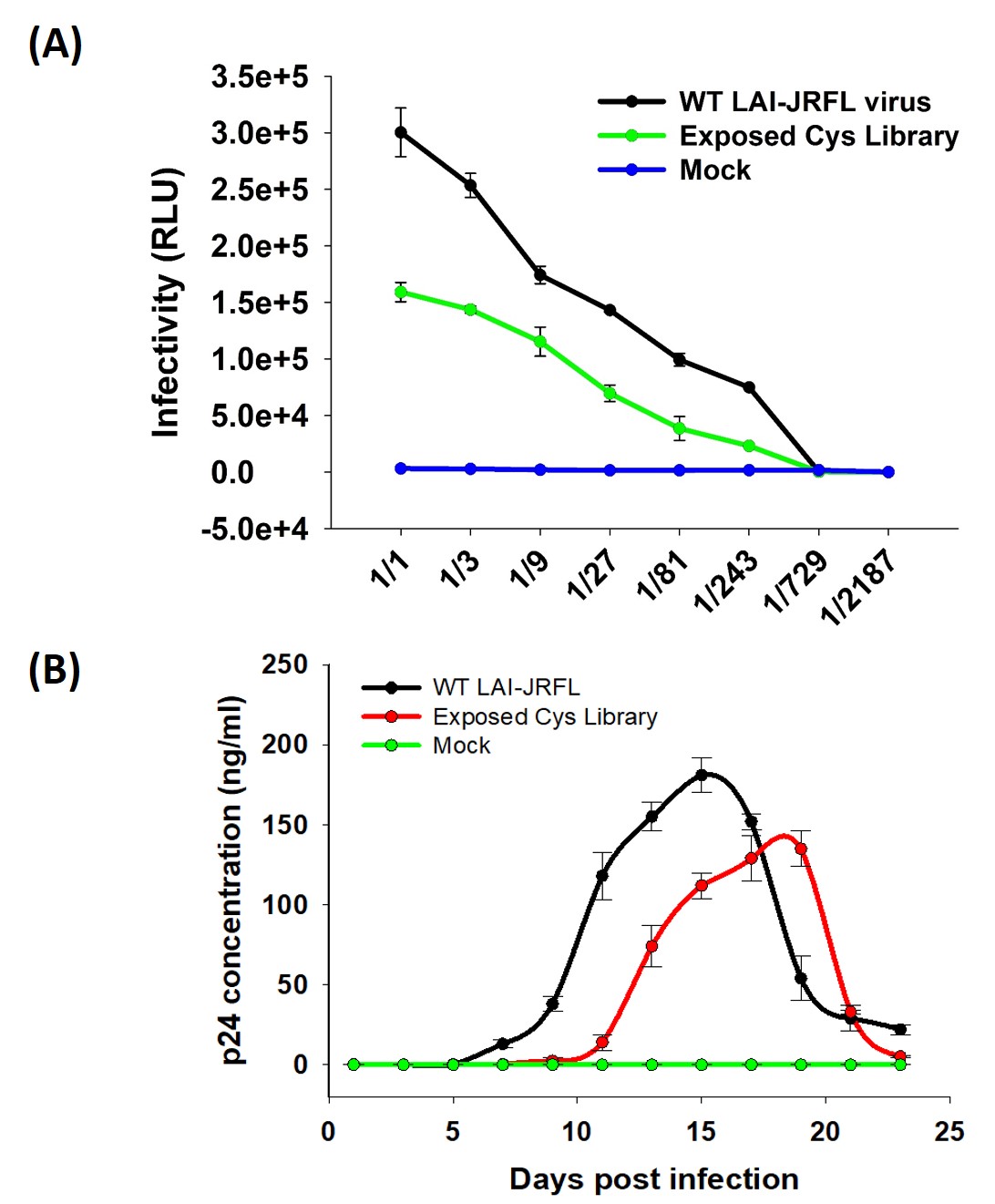

Fig. S3. Infectivity of ExCysLib in comparison to WT virus (A) Single-cycle HIV-1 infectivity in TZM-bl cells and (B) multi-cycle HIV-1 infectivity in HUT-R5 cells.

(A) For single-cycle infectivity assays, Relative luminescence units (RLUs) were plotted as a function of viral dilution. Equivalent p24 amounts were used for WT and Exposed Cys Library (ExCysLib)

(B) For multi-cycle infectivity assays, p24 levels were estimated using the HIV-1 p24 SimpleStep ELISA kit (Abcam). The ExCysLib was found to retain significant viral infectivity in both single-cycle and multi-cycle viral infectivity assays. Equivalent p24 amounts were used for initial infection. Error bars represent SDs of two replicate experiments.

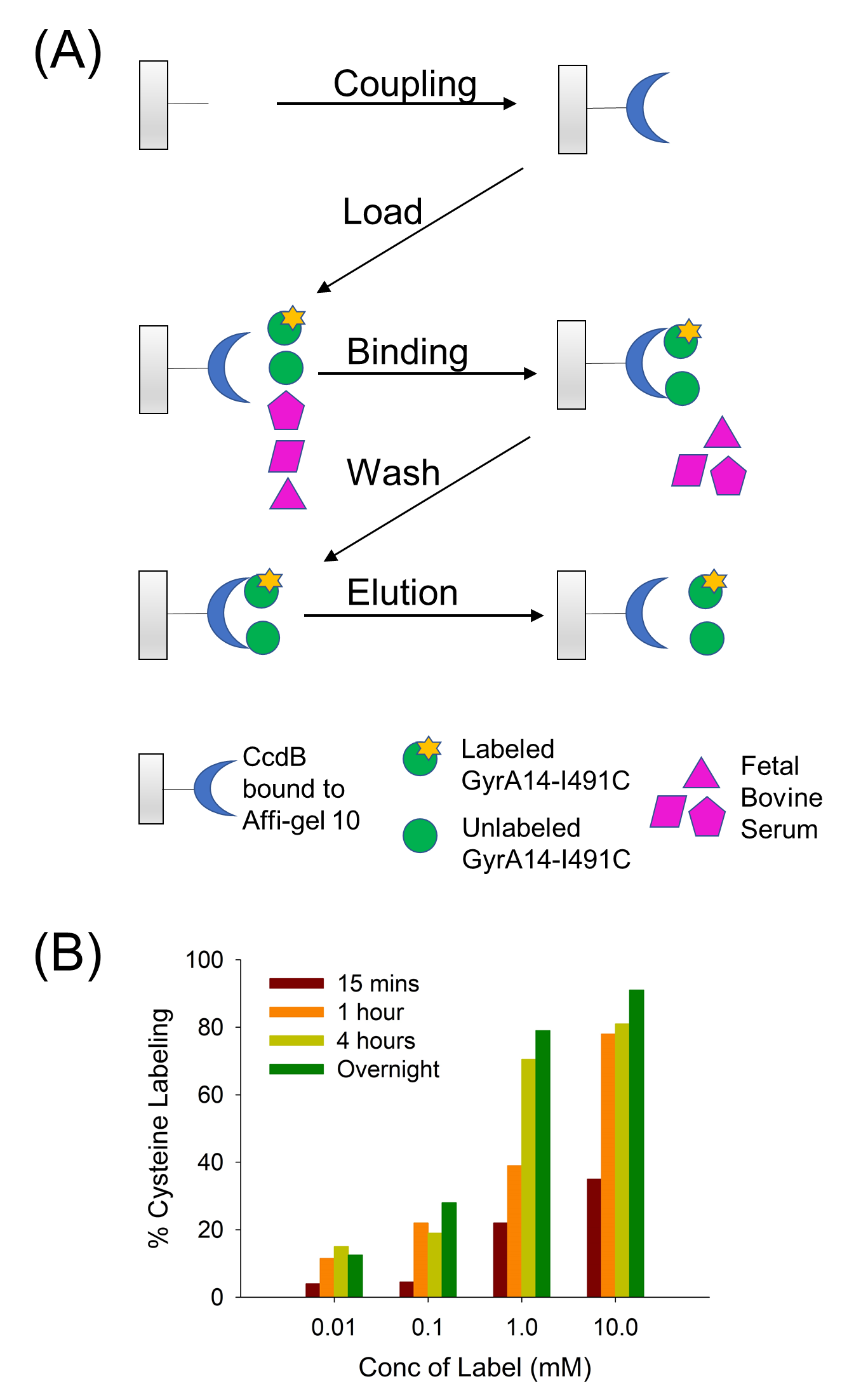

Fig. S4. Quantification of the extent of Cys labeling, using labeled GyrA14-I491C as a test protein.

(A) The extent of incorporation of the Cys label, Maleimide-PEG2-Biotin, was quantified using GyrA14 (residues 363-494 of DNA Gyrase) with a Cys mutation at a solvent-exposed residue I491 (GyrA14-I491C) as a test protein. Labeling of test protein was carried out in 20% FBS to mimic viral labeling in the presence of FBS. Subsequently, affinity purification of the labeled GyrA14-I491C protein was carried out using a CcdB-immobilized column. Both the labeled and unlabeled test protein bind to the CcdB column.

(B) Elutes were subsequently analyzed by MALDI-TOF mass spectrometry. The relative peak intensities of the labeled and unlabeled proteins were used to estimate the extent of Cys labeling.

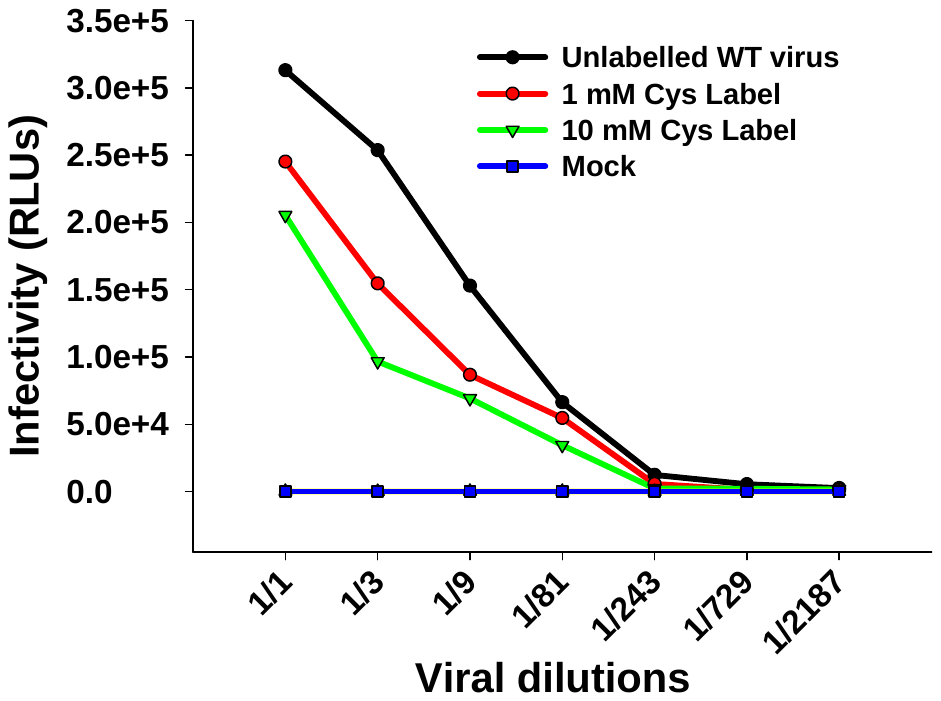

Fig. S5. Single-cycle infectivity assay of the WT virus in the presence of 1 mM and 10 mM of Cys label, Maleimide-PEG2-Biotin, in TZM-bl cells.

Relative luminescence units (RLUs) were plotted as a function of viral dilution. The WT virus retained significant infectivity in the presence of 10 mM label, in comparison to the unlabeled virus. Labeling was carried out in 1X PBS, overnight at 4°C.

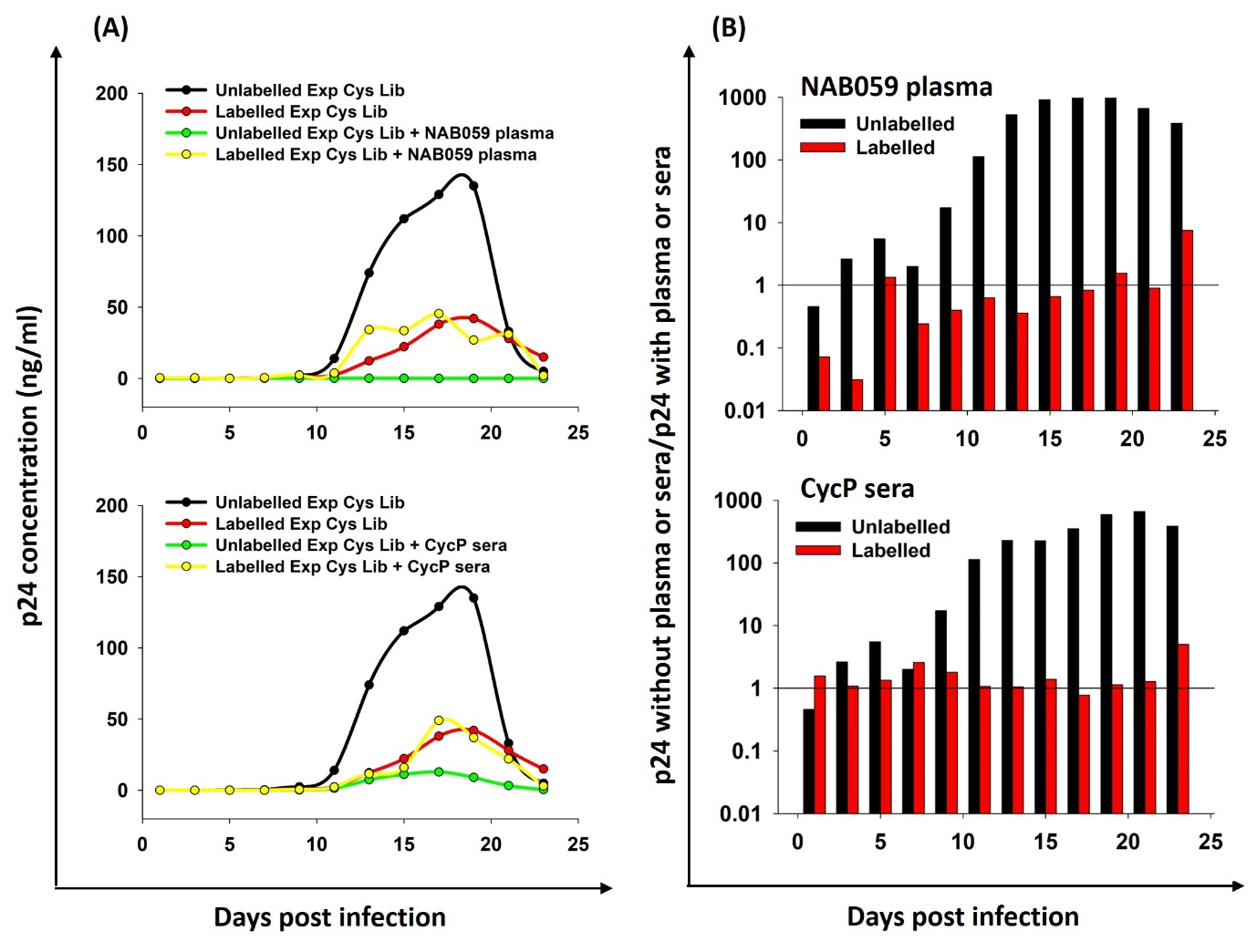

Fig. S6. Multi-cycle infectivity of the ExCysLib in the presence of Cys labeling reagent as well as NAB059 plasma (top) or trimeric cyclic permutant (CycP)-immunized guinea pig sera (bottom) in HUT-R5 cells.

The ratio of p24 levels for the labeled ExCysLib in the absence to presence of NAB059 plasma and CycP sera was quantified and was found to be ~1 (red bar), confirming that labeling of one or more Cys residues present in the ExCysLib makes the virus resistant to neutralization. Infectivity ratios <1 at initial time points are attributed to very low levels of virus and are likely artifactual.

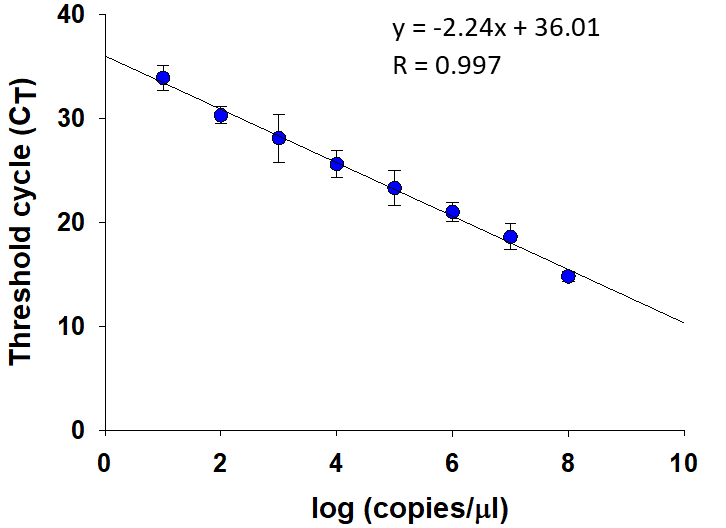

Fig. S7. Standard curve for viral titer estimation using qRT-PCR.

The standard curve was plotted using threshold cycle (C_T_) values obtained from qRT-PCR data of the pol gene amplified from different dilutions of the pLAI-JRFL plasmid ranging from 10^1^ to 10^8^ copies/µl. SYBR green detection chemistry was used for qRT-PCR. The standard curve was used for absolute quantitation of viral cDNA. Error bars represent SDs of two replicate experiments.

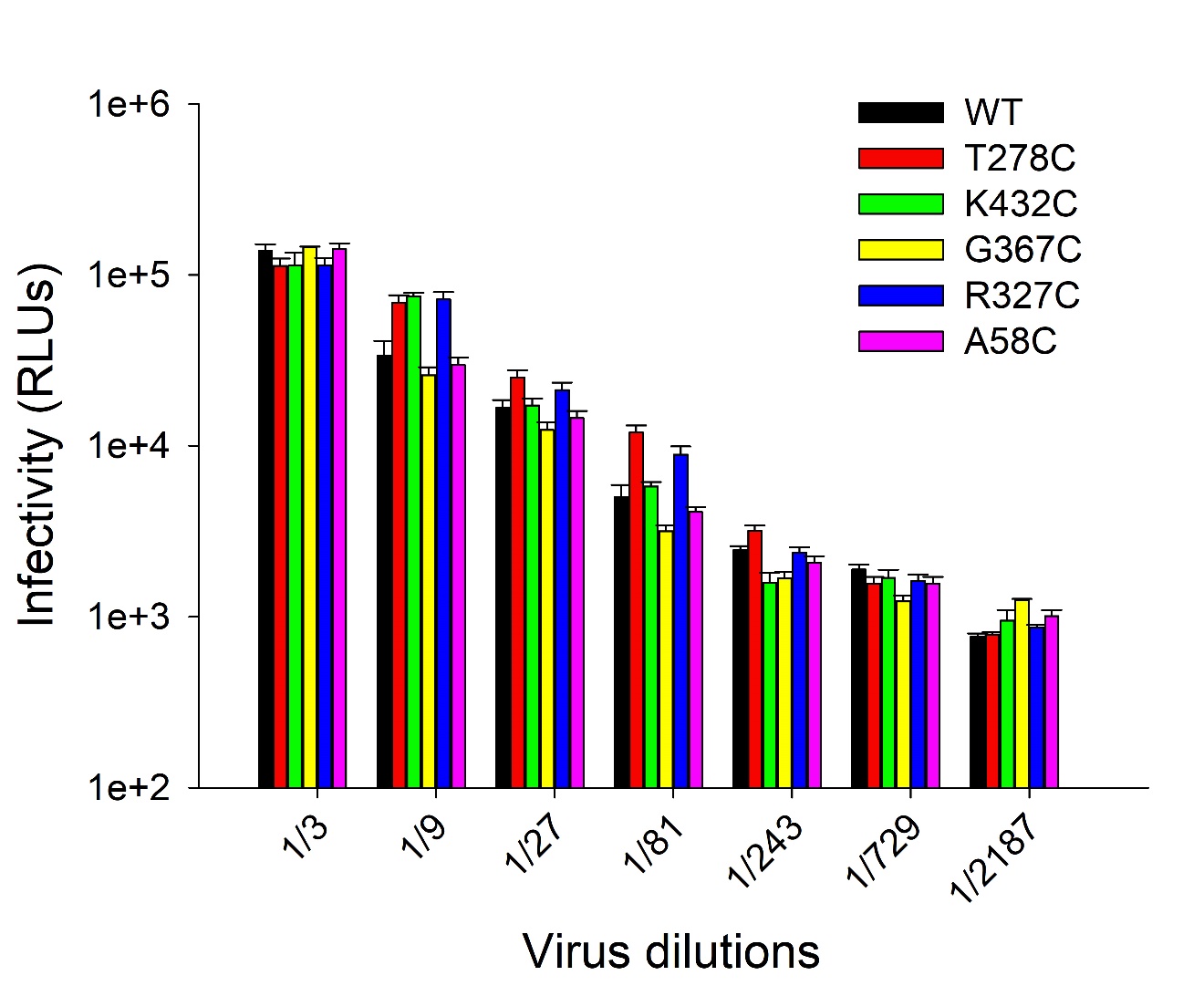

Fig. S8. Single-cycle infectivity of bNAb epitope Cys mutant viruses in TZM-bl cells.

The single-cycle infectivity of the WT virus, as well as Cys mutant viruses in TZM-bl cells. Infection was monitored by measuring the relative luminescence units (RLUs) which are plotted above as a function of the viral dilution. All epitope-Cys mutant viruses were found to be similarly infectious as the WT virus in TZM-bl cells, at all viral dilutions examined. Equivalent amounts of p24 were taken for all viruses. Error bars represent SDs of two replicate experiments.

Table S1. List of exposed residues in Env selected for Cys mutagenesis.

| **#** | **Residue Number^§^** | **Residue in LAI-JRFL** | **Residue in BG505** | **Solvent Accessibility^¥^ (%)** | **Distance between side-chain centroids^ψ^ from nearest Cys mutant in ExCysLib (Å)** |
| --- | --- | --- | --- | --- | --- |
| 1 | 32 | E | E | 64.4 | 8.1 (from 500) |
| 2 | 49 | T | E | 52.2 | 11.7 (from 106) |
| 3 | 58 | A | A | 42.1 | 10.7 (from 75) |
| 4 | 63 | T | T | 36.7 | 9.1 (from 211) |
| 5 | 66 | H | H | 52.4 | 12.0 (from 63) |
| 6 | 73 | A | A | 42.8 | 8.5 (from 75) |
| 7 | 75 | V | V | 32.7 | 8.5 (from 73) |
| 8 | 78 | D | D | 46.3 | 9.9 (from 75) |
| 9 | 83 | E | E | 33.8 | 12.9 (from 78) |
| 10 | 88 | N | N | 53.4 | 9.1 (from 90) |
| 11 | 90 | T | T | 45.3 | 9.1 (from 88) |
| 12 | 94 | N | N | 31.9 | 8.0 (from 275) |
| 13 | 106 | E | T | 37.2 | 11.7 (from 49) |
| 14 | 117 | K | K | 56.8 | 9.1 (from 432) |
| 15 | 121 | K | K | 43.8 | 10.3 (from 117) |
| 16 | 132 | K | T | 40.2 | 8.3 (from 188) |
| 17 | 135 | N | T | 66.2 | 8.7 (from 132) |
| 18 | 137 | T | N | 60.6 | 10.2 (from 135) |
| 19 | 140 | T | D | 67.2 | 10.0 (from 151) |
| 20 | 151 | R | R | 39 | 10.0 (from 140) |
| 21 | 160 | N | N | 30.1 | 12.1 (from 168) |
| 22 | 166 | R | R | 43.5 | 12.8 (from 168) |
| 23 | 168 | E | K | 60.1 | 12.1 (from 160) |
| 24 | 183 | P | Q | 49.8 | 8.1 (from 185) |
| 25 | 185 | N | N | 86.2 | 8.1 (from 183) |
| 26 | 187^ | N | S | 100 | 6.1 (from 188) |
| 27 | 188 | N | N | 47.6 | 6.1 (from 187) |
| 28 | 194 | I | I | 31.6 | 11.8 (from 183) |
| 29 | 211 | E | E | 47.1 | 9.1 (from 63) |
| 30 | 232 | T | K | 56 | 9.7 (from 267) |
| 31 | 234 | N | N | 36.8 | 9.7 (from 232) |
| 32 | 267 | E | E | 32.6 | 9.7 (from 232) |
| 33 | 275 | D | E | 30.2 | 8.0 (from 94) |
| 34 | 278 | T | T | 58.6 | 10.5 (from 281) |
| 35 | 281 | A | A | 47.5 | 9.7 (from 275) |
| 36 | 293 | E | Q | 37.4 | 11.0 (from 444) |
| 37 | 306 | S | S | 42.4 | 10.3 (from 440) |
| 38 | 314 | G | G | 37.9 | 10.7 (from 121) |
| 39 | 323 | I | I | 39.1 | 10.3 (from 442) |
| 40 | 325 | D | D | 58.1 | 10.4 (from 323) |
| 41 | 327 | R | R | 43.5 | 8.6 (from 442) |
| 42 | 335 | R | K | 46.4 | 7.8 (from 412) |
| 43 | 340 | D | E | 56.9 | 10.9 (from 335) |
| 44 | 347 | I | K | 37.5 | 8.1 (from 351) |
| 45 | 351 | E | K | 44.2 | 8.1 (from 347) |
| 46 | 354 | E | G | 35.4 | 8.5 (from 463) |
| 47 | 365 | S | S | 58.2 | 8.1 (from 367) |
| 48 | 367 | G | G | 54.7 | 8.1 (from 365) |
| 49 | 386 | N | N | 30.5 | 10.8 (from 392) |
| 50 | 392 | N | N | 57.9 | 8.6 (from 394) |
| 51 | 394 | T | T | 42.3 | 8.6 (from 392) |
| 52 | 398 | N | N | 82.2 | 9.4 (from 347) |
| 53 | 412 | N | N | 84.9 | 8.1 (from 335) |
| 54 | 415 | T | T | 43.3 | 8.3 (from 412) |
| 55 | 428 | Q | Q | 74.3 | 12.1 (from 367) |
| 56 | 432 | K | Q | 31.9 | 9.1 (from 117) |
| 57 | 440 | R | Q | 41.3 | 10.3 (from 306) |
| 58 | 442 | Q | V | 46.3 | 8.2 (from 444) |
| 59 | 444 | R | R | 61.7 | 8.2 (from 442) |
| 60 | 459 | G | G | 58.1 | 11.4 (from 365) |
| 61 | 463 | N | S | 68.9 | 8.5 (from 354) |
| 62 | 500 | K | R | 33.9 | 8.1 (from 32) |
| 63 | 505 | V | V | 88.1 | 10.0 (from 658) |
| 64 | 518 | V | V | 86.1 | 9.8 (from 535) |
| 65 | 535 | M | M | 42.6 | 9.8 (from 518) |
| 66 | 547 | G | G | 48.8 | 12.6 (from 75) |
| 67 | 569 | T | T | 82 | 10.9 (from 117) |
| 68 | 609 | P | P | 46.3 | 8.1 (from 645) |
| 69 | 612 | A | S | 45.2 | 10.2 (from 609) |
| 70 | 617 | K | R | 33.9 | 8.4 (from 625) |
| 71 | 620 | D | S | 42.4 | 8.2 (from 625) |
| 72 | 625 | N | N | 55.2 | 8.0 (from 630) |
| 73 | 630 | E | Q | 39.7 | 8.0 (from 625) |
| 74 | 633 | R | K | 53.4 | 8.5 (from 630) |
| 75 | 637 | N | N | 56.6 | 9.2 (from 633) |
| 76 | 640 | S | Q | 56.4 | 11.0 (from 645) |
| 77 | 645 | L | L | 30.1 | 8.1 (from 609) |
| 78 | 653 | Q | Q | 46.7 | 10.8 (from 660) |
| 79 | 658 | Q | Q | 33.1 | 9.8 (from 660) |
| 80 | 660 | L | L | 39.6 | 8.5 (from 664) |
| 81 | 664 | D | D | 68.1 | 8.5 (from 660) |

^§^Residues are numbered according to the HxB2 numbering scheme (<https://www.hiv.lanl.gov/content/sequence/HIV/REVIEWS/HXB2.html>)

^¥^Solvent accessibility of each residue was calculated using the Naccess V 2.1.1 program for the BG505 Env (PDB ID: 4TVP)

**^ψ^**Pairwise side-chain side-chain centroid distances were calculated using in-house PERL scripts for each of the indicated bNAbs

^Residue 187 was included even though it is in close proximity to residue 188, on account of it having very high (100%) solvent accessibility.

Table S2. Neutralization Potency (IC_50_) of bNAbs for HIV-1 LAI-JRFL virus.

| **bNAb** | **Binding Site on Env** | **Reported IC_50_ with JRFL (ng/ml) (Reference)** | **Measured IC_50_ with LAI-JRFL (ng/ml)** |
| --- | --- | --- | --- |
| VRC01 | CD4bs | 31 (1) | 41 ± 2.8 |
| b12 | CD4bs | 22 (1) | 43 ± 2.8 |
| PGT128 | V3-glycan | 7 (2) | 12 ± 3.3 |
| PGT151 | gp120-gp41 interface | 10 (Clade B) (3) | 28 ± 3.2 |

Table S3. List of MID sequences for each condition used in deep sequencing.

| MID # | Condition | MID Sequence |
| --- | --- | --- |
| 1 | Labeled ExCysLib + bNAb VRC01 | TGGTCA |
| 2 | Labeled ExCysLib + bNAb PGT128 | CGATGT |
| 3 | Labeled ExCysLib + bNAb PGT151 | TGACCA |
| 4 | Labeled ExCysLib + NAB059 plasma | GCCAAT |
| 5 | Labeled ExCysLib + CycP sera | CAGATC |
| 6 | Labeled ExCysLib, no antibody | CTTGTA |
| 7 | Unlabeled ExCysLib, no antibody | AGTCAA |
| 8 | ExCysLib mutant plasmid DNA library | AGTTCC |

Table S4. bNAb epitope residues identified from deep sequencing data.

| **VRC01 bNAb**  **(MID 1)** | | **PGT128 bNAb (MID 2)** | | **PGT151 bNAb (MID 3)** | | **NAB059 plasma (MID 4)** | **CycP sera (MID 5)** |
| --- | --- | --- | --- | --- | --- | --- | --- |
| Identified | Known^Ø^ | Identified | Known^Ø^ | Identified | Known^Ø^ | Identified | Identified |
| T278 | T278 | - | N135^¥^ | A58 | A58 | A73 | T140 |
| A281 | A281 | I323 | I323 | T63^§^ | - | S365 | R183 |
| S365 | S365 | D325 | D325 | - | V518^¥^ | N412 | I194 |
| G367 | G367 | R327 | R327 | G547 | G547 | N463 | Q428 |
| Q428 | Q428 | R444 | R444 | R633^§^ | - | N625 | K432 |
| G459 | G459 | **-** | **-** | N637 | N637 | - |  |
| N463 | N463 | **-** | **-** | **-** | S640^¥^ | **-** | **-** |

^Ø^Structurally defined epitopes are indicated for each bNAb

^¥^Structurally defined epitope, not identified by us (False negative)

^§^Not part of structurally defined epitope, but identified by us (possible false positive)

Table S5. Individual epitope residues for each bNAb selected for validation studies using Cys labeling.

| **Epitope residue** | **bNAb** | **Epitope on Env** | **PDB ID** |
| --- | --- | --- | --- |
| T278 | VRC01 | CD4-binding site | 3NGB^§^ |
| K432 | b12 | CD4-binding site | 2NY7^§^ |
| G367 | VRC01, b12 | CD4-binding site | 3NGB^§^, 2NY7^§^ |
| R327 | PGT128 | V3-glycan | 5ACO^§^ |
| A58 | PGT151 | gp120-gp41 interface | 5FUU^¥^ |

Epitopes of bNAbs on Env were identified from available X-ray^§^ or cryo-EM^¥^ structural information.

**Table S6. Viral titer estimation of WT and Cys-mutant viruses.** Viral titers were estimated by absolute quantitation of viral cDNA from the standard curve, obtained after qRT-PCR of the pol gene.

| **Epitope mutant** | **bNAb** | **Threshold cycle (C_T_)** | **Titer**  **(x 10^5^ copies/µl)** |
| --- | --- | --- | --- |
| WT | - | 23.1 | 1.06 |
| T278C | VRC01 | 24.4 | 0.34 |
| K432C | b12 | 24.8 | 0.24 |
| G367C | VRC01, b12 | 22.2 | 2.5 |
| R327C | PGT128 | 23.9 | 0.53 |
| A58C | PGT151 | 25.4 | 0.14 |
